# CD133⁺ Vesicles Mediate Resistance to RAS-ERK Inhibition Regulated by YAP Activation in Liver Cancer Cells

**DOI:** 10.1101/2025.07.31.667712

**Authors:** Shuo Zhang, Xue Feng, Evelyn Quan, Kota Kaneko, Gen-Sheng Feng

## Abstract

CD133, a pentaspan plasma membrane protein, has been viewed as a biomarker of stem cells in normal and cancer tissues, although its function and mechanism are unclear. In previous studies, we identified a new type of CD133^+^ intracellular vesicles, named intercellsome, which is implicated in direct cell-cell communication under stress conditions. However, the regulatory mechanism and biological significance of these CD133^+^ vesicles are largely unknown, highlighting a gap in understanding of this cellular communication mechanism. We show here that CD133 acts as a stress response marker in cancer cells, with its expression and vesicle formation significantly induced under MEK inhibitor-mediated proliferative stress. The CD133^+^ vesicles are essential for maintaining cell proliferation under stress conditions *in vitro*. Mechanistically, the MEKi activates the Hippo-YAP signaling pathway, which promotes CD133 transcription, establishing a novel connection between YAP signaling and CD133^+^ vesicle biogenesis. Further, CD133 plays a critical role in YAP-driven liver cancer progression in mice. This study defines a critical role of CD133^+^ vesicles in stress response regulated by YAP, which advances the understanding of CD133 functions beyond its stem cell-associated roles and suggests new avenues for therapeutic intervention of liver cancer relapse.

## Introduction

A major challenge in cancer therapy is the drug resistance, where cancer cells survive and continue to proliferate despite the treatment (1–3). This resistance often involves proliferative defects that enable cancer cells to adapt to stress and evade cell death, posing a significant barrier to successful clinical outcomes (2, 3). The SHP2-RAS-ERK signaling pathway is a well-characterized cascade that regulates cell proliferation, survival and differentiation (4, 5). Active receptor tyrosine kinases (RTKs) such as EGFR and MET could activate the SHP2 phosphatase activity, which dephosphorylates RasGAP and to promote RAS activation (5, 6). SHP2 is required for full activation of the RTK/RAS/ERK signaling. Loss of SHP2 leads to proliferation defects, as evidenced by impaired hepatocyte proliferation following partial hepatectomy and suppressed RTK-driven liver cancer progression (5, 7, 8).

CD133 (also known as AC133 or Prominin-1) is a pentaspanin transmembrane glycoprotein. CD133 signal was located in the lipid raft regions of plasma membrane (9), and was enriched in endosome in pericentrosomal region (10). CD133 has been implicated in several signal pathways, such as Wnt/β-Catenin, IL-6, TGF-β, AKT and mTOR (11–16). Of note, this cell surface protein has been viewed as a biomarker of cancer stem cells (CSC) in cancers of brain, lung, colon, liver, ovarian and pancreatic origins (16–19). CD133^+^ CSCs demonstrate stronger resistance to chemotherapy compared to their CD133^-^ counterparts (20–23). Elevated CD133 expression levels are associated tumor recurrence and poor prognosis. However, CD133 negative tumor cells were also shown to form clonospheres *in vitro* and tumors in immune-deficient mice *in vivo*. Regardless of the controversies, the function and mechanism of CD133 remain to be determined.

In previous experiments aimed at dissecting hepatocyte proliferation during liver regeneration, we identified a new type of intracellular vesicles, named intercellsomes (7). These vesicles are positive for CD133, a pentaspanin, which are distinctive from the classical exosomes that are labeled by tetraspanins, such as CD9, CD63 and CD81 (7). Notably, CD133- positive vesicles were predominantly induced in Shp2-deficient hepatocytes during liver regeneration, following partial hepatectomy or CCl_4_-triggered liver injury. The CD133-positive vesicles contained mitogenic mRNAs rather than miRNAs that are enriched in exosomes. Our preliminary data also suggest that the CD133-positive vesicles migrate between tightly contacting hepatocytes, which may mediate sharing of the mitogenic mRNAs and enable cell proliferation. Therefore, we proposed that formation of the CD133^+^ vesicles is a stress responsive mechanism activated under intracellular proliferative signal deficit. However, many open questions remain to be answered to establish this theory. In particular, the precise function of CD133⁺ vesicles and their regulatory mechanisms are largely unknown.

Yes-associated protein (YAP) is a key player of the Hippo signaling pathway in regulating cell proliferation, tissue homeostasis and cancer development (24–30). Typically, upstream MST1/2 kinases phosphorylate and activate LATS1/2 kinases, which in turn phosphorylate YAP and its homolog TAZ at multiple sites. Phosphorylated YAP and TAZ are retained in the cytoplasm and prevented from entering the nucleus, thereby inhibiting their transcriptional activity (25, 27–29, 31). The YAP/TAZ activity is finely tuned by a complex network of intracellular and intercellular signaling pathways (27–29). Elevated YAP activity has been associated with enhanced cell proliferation, wound repair, tissue regeneration, and tumorigenesis (29). However, it remains to be determined if and how YAP activity influences CD133 expression and formation of CD133^+^ vesicles in the context of cell proliferation under stress.

In this study, we systematically screened multiple cancer cell lines and isolated CD133^+^ vesicles to elucidate their unique properties. A comparative RNA cargo analysis demonstrated distinct features of CD133^+^ vesicles, as compared to extracellular vesicles (EVs). Moreover, we unveiled a critical role of YAP in regulation of CD133 expression and CD133⁺ vesicles in driving cell proliferation and tumorigenesis, highlighting a novel pathway that may contribute to cancer relapse and therapeutic resistance.

## Results

### CD133 is upregulated in response to proliferation stress in multiple cancer cell lines

To investigate the properties of CD133^+^ vesicles, we first screened multiple cell lines to assess the basal levels of CD133 expression. Consistent with previous reports, we observed that CD133 expression is particularly higher in liver cancer cells and the colon cancer cell line CaCo2 (Supplementary Figure 1A). Additionally, we also examined potential correlations between CD133 and other tetraspanin proteins (CD9, CD63, and CD81), commonly recognized as extracellular vesicle (EV) markers (Supplementary Figure 1A). Notably, CD133 expression level did not significantly correlate with the expression pattern of CD63, CD9 and CD81, suggesting that CD133 may not be associated with canonical EV populations. To establish a model for proliferation defects *in vitro*, we treated PLC and Huh7 cell lines with MEKi. We found that MEKi treatment significantly reduced cell proliferation without inducing cytotoxicity (Supplementary Figure 1B-1C), effectively recapitulating the proliferation defect observed in the partial hepatectomy model of liver-specific Shp2 knockout mice (7). These results confirm the reliability of this *in vitro* proliferation defect model.

Then, we examined CD133 expression level changes in PLC and Huh7 cells treated with SHP2i and MEKi. Both of SHP2i and MEKi treatments significantly increased CD133 at both mRNA and protein levels (Figure 1A-1D). To minimize background signaling from serum-derived factors, we performed serum starvation prior to treatment, and found CD133 induction remained evident even under serum-starved conditions. (Supplementary Figure 1D). We further evaluated whether CD133 upregulation in response to SHP2i and MEKi is a generalizable phenomenon across various cell lines. Consistently, all tested cell lines exhibited significant induction of CD133 expression following inhibitor treatment (Supplementary Figure 2A, 2B), suggesting that CD133 upregulation is a common cellular response to proliferative defects. To further rule out off-target effects of individual inhibitors, we utilized two additional MEKi and one ERKi. All treatments resulted in a significant increase in CD133 expression at both the mRNA and protein levels, reinforcing the conclusion that CD133 induction is a consistent cellular response to impaired proliferation (Supplementary Figure 3A-3C).

**Figure 1.**
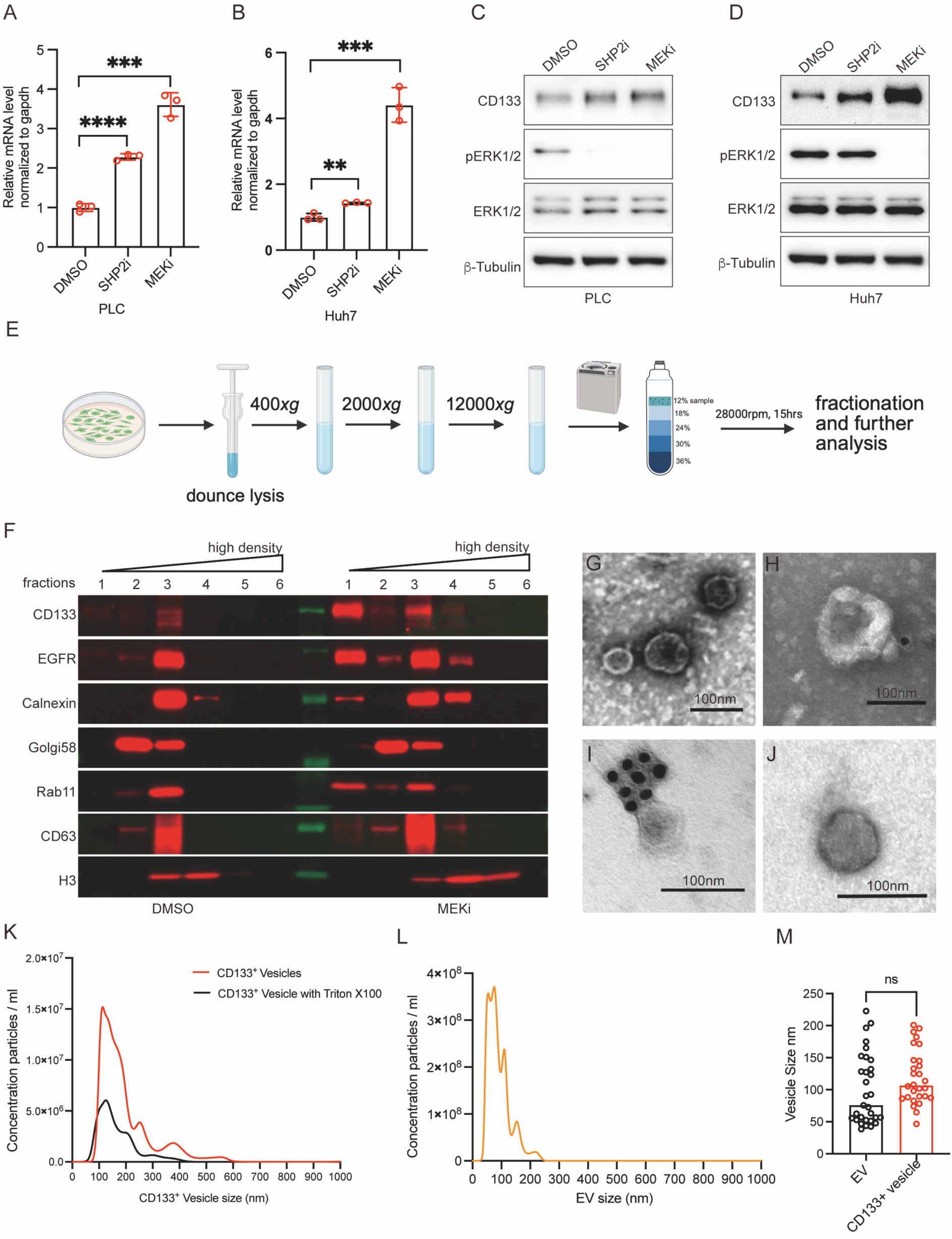
CD133^+^ vesicle is upregulated in response to SHP2i and MEKi induced proliferation impairment. (A-B) mRNA level of CD133 were analyzed when treating cells with SHP2i or MEKi for 48 hours. (A) PLC cells, (B) Huh7 cells. Statistical significance is determined by Student’s t-test. Data were represented as mean ± SEM. *p<0.05, **p<0.01, ***p<0.001. (C-D) Protein levels of CD133, pERK1/2 and ERK1/2 were analyzed by western blotting when cells were treated with SHP2i or MEKi for 48 hours. There is a significant upregulation of CD133 after inhibitor treatment. (C) PLC cells, (D) Huh7 cells. (E) Diagram of the vesicle isolation and purification system. Cells were lysed using a Dounce homogenizer and the lysate was processed by differential centrifugation. The resulting supernatant was subjected to high- resolution density gradient ultracentrifugation followed by fractionation. The top fraction, enriched in vesicles, was collected for further analysis. (F) PLC fractions from DMSO control and MEKi treatment were analyzed by western blotting. Fraction 1 to 6 goes from low density to high density. Same cell number were used in two conditions. CD133 is upregulated in the top fractions. EGFR is the plasma membrane marker, Calnexin is an ER marker, Golgi58 is a Golgi marker. (G) Representative negative staining electron microscope picture of fraction 1 in PLC MEKi condition, scale bar, 100nm. (H-I) Representative immune-gold labeling electron microscope picture of fraction 1 in PLC MEKi condition, gold particle size is 18nm, scale bar, 100nm. (J) Representative negative staining electron microscope of EV from PLC supernatant, scale bar, 100nm. (K) NTA of indicated CD133^+^ vesicles from MEKi treated PLC cells showing size distribution (n=3 biological replicates). Vesicles with same amount were treated for 5% Triton X-100 for 30 minutes at room temperature. (H) NTA of indicated EV from PLC supernatant showing size distribution (n=3 biological replicates). (I) CD133^+^ vesicle size and EV size comparation based on the immune-gold labeling electron microscope image, each dot represents a vesicle. Vesicle size was calculated by ImageJ (NIH). Statistical significance is determined by Student’s t-test. ns: no significance.

### MEKi induces CD133^+^ vesicles in liver cancer cells that are distinct from EVs

Given the significant upregulation of CD133 following MEKi treatment, we sought to determine whether these upregulated CD133 molecules were associated with vesicles. In order to isolate and detect CD133^+^ vesicles, we first set up the vesicle isolation and purification system combining the differential centrifugation with high-resolution iodixanol density gradient fractionation method (Figure 1E). Cells were lysed using a dounce homogenizer and cell lysate underwent differential centrifugation to collect the supernatant, which was then subjected to high-resolution density gradient ultracentrifugation and fractionation. Individual fractions were collected, and protein contents were analyzed (Figure 1E). As previously reported (32), the top fractions primarily contained small vesicles. Enrichment of CD133^+^ vesicles, but not CD63^+^ vesicles, was detected in the top fraction following MEKi treatment compared to the DMSO control, indicating that MEKi-induced CD133^+^ vesicles may represent a specific class of small vesicles (Figure 1F). This enrichment of CD133^+^ vesicles in the top fraction was also confirmed in vesicle fractions derived from Huh7 cells (Supplementary Figure 4). Treatment of the cell lysate with detergent led to the loss of membrane protein signals after ultracentrifugation, highlighting that CD133 is predominantly membrane-associated (Supplementary Figure 5A, 5B). Furthermore, we adopted the electron microscopy to examine the top fraction via negative staining, and observed obvious vesicular structures (Figure 1G). Subsequent immune-gold labeling further confirmed the presence of CD133^+^ vesicles in the top fraction (Figure 1H, 1I), which comprise approximately 5% of the total vesicle population (data not shown). We also assessed EV morphology from the supernatant of PLC cells (Figure 1J) and found the size of EV was comparable to that of CD133^+^ vesicles (Figure 1K-1M). Treatment of the purified CD133^+^ vesicles with detergent caused a significant decrease in vesicle concentration, further confirming their vesicle structures (Figure 1K). To investigate whether CD133^+^ vesicles are related to EVs, cells were treated with GW4869, an EV biogenesis inhibitor (33, 34). This treatment increased CD63 levels but did not affect CD133, suggesting that the induction of CD133^+^ vesicles by MEKi is independent of EV production (Supplementary Figure 5C).

### Transcriptomic analysis reveals unique RNA signatures in CD133^+^ vesicles

Given the established role of EVs in cell-cell communication through their cargo (32–34), we investigated whether CD133^+^ vesicles carry cargo and whether their cargo profiles resemble those of EVs. We employed density fractionation combined with immunoprecipitation to isolate pure CD133^+^ vesicles, and subsequently analyzed the RNA content of CD133^+^ vesicles alongside that of total cells and EVs. Principal component analysis (PCA) of the total RNA expression profiles revealed a clear separation among total cells, CD133^+^ vesicles, and EVs, with EVs clustering more closely with total cells (Supplementary Figure 6A). Volcano plots of CD133^+^ vesicles with total cells and with EVs both revealed significant differential gene expression, further confirming the distinct RNA content between CD133^+^ vesicles and EVs (Supplementary Figure 6B-6D). In contrast, small RNAs, particularly miRNAs, exhibited more distinct difference among them (Supplementary Figure 6E). Moreover, gene expression heat map analysis showed that EVs and CD133^+^ vesicles have distinct enriched gene sets (Supplementary Figure 6F), demonstrating that they represent two distinct classes of vesicles.

### CD133^+^ vesicles promote cell proliferation *in vitro*

To investigate the effect of CD133^+^ vesicles on cell growth *in vitro*, we treated cells with vesicles derived from various sources and assessed cell viability afterward (Figure 2A). To verify vesicle uptake by recipient cells, CaCo2 cells were incubated with PKH26-labeled vesicles derived from PLC cells (Figure 2B). The detection of PKH26 signals within the cells confirmed active vesicle uptake. Next, we isolated CD133^+^ vesicles enriched vesicle fraction from MEKi treated PLC cells and applied them to various recipient cell lines, including HeLa, 293T, and PLC cells (Figure 2C-2E). All recipient cell lines exhibited a significant increase in viability compared to controls, demonstrating that CD133^+^ vesicles enhance cell proliferation. Additionally, we compared the effects of vesicles derived from PLC and CaCo2 cells, which differ in CD133 levels (Supplementary Figure 1A). Vesicles from CaCo2 cells, characterized by higher CD133 content (Supplementary Figure 5D, 5E), exhibited a more pronounced proliferation-promoting effect than those from PLC cells, suggesting a dosage-dependent relationship between CD133 content and proliferation enhancement (Figure 2C).

**Figure 2.**
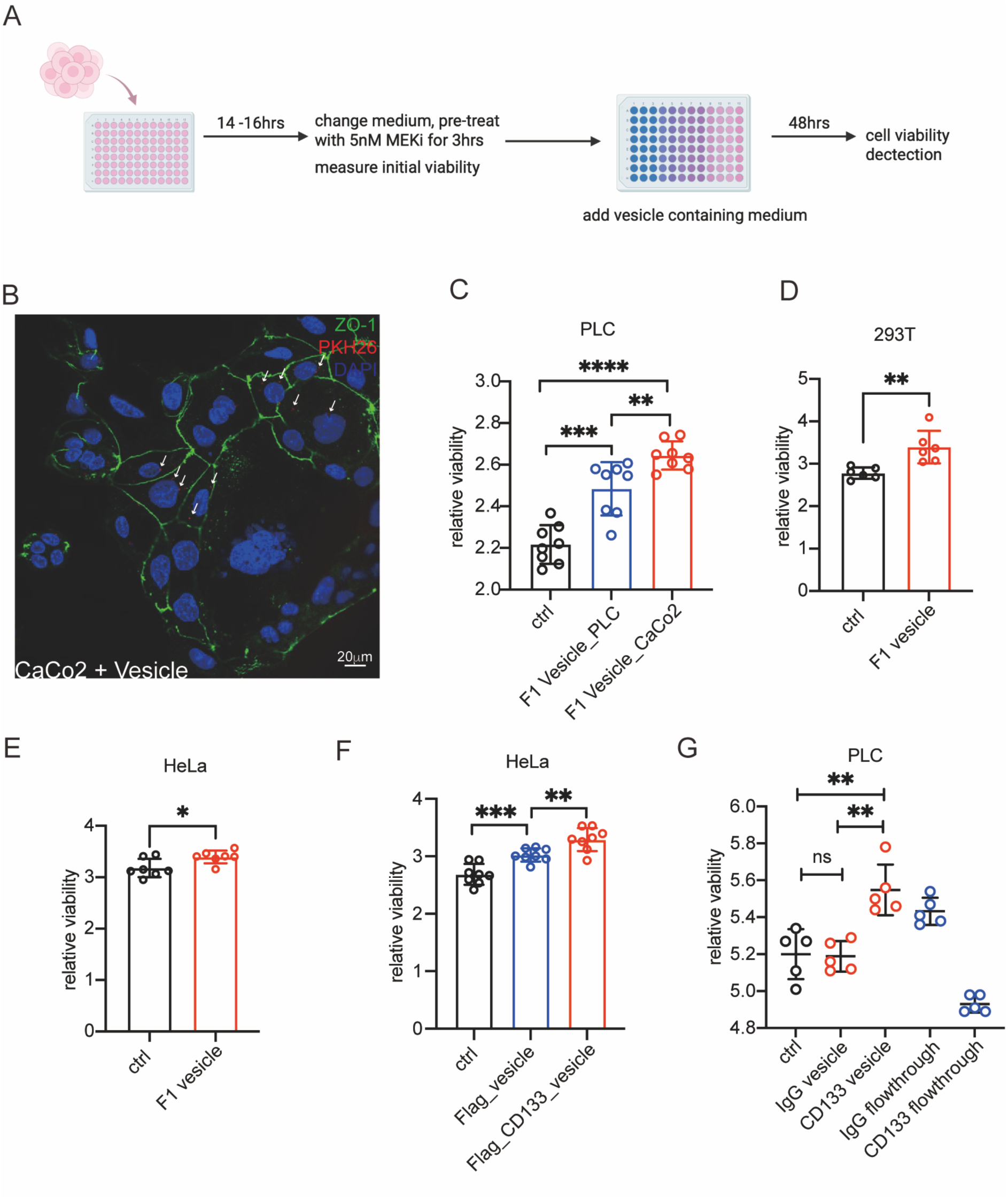
CD133^+^ vesicles promote cell proliferation *in vitro*. (A) Schematic of the cell proliferation assay. Cells were seeded into 96-well plates one day before treatment. MEKi pre-treatment was performed, followed by the addition of vesicles. After 48 hours of incubation, cell viability was assessed using the CCK-8 assay. (B) Top fraction enriched with CD133⁺ vesicles was stained with PKH26 to visualize vesicle uptake. CaCo2 cells were used as recipient cells. Cells were fixed 3 hours after vesicle treatment. ZO-1 (green), PKH26 (red), DAPI (Blue). Scale bar, 20 μm. (C) 293T cells were treated with CD133^+^ vesicles enriched fractions; cell viability was determined after 48 hours of incubation. (D) HeLa cells were treated with CD133^+^ vesicles enriched fractions; cell viability was determined after 48 hours of incubation. (E) PLC cells were treated with CD133^+^ vesicles enriched fractions from both PLC cells and CaCo2 cells with same number, CaCo2 cells have higher CD133 levels than PLC cells. Cell viability was determined after 48 hours of incubation. (F) HeLa cells were treated with CD133⁺ vesicle-enriched fractions isolated from PLC cells stably expressing either Flag or Flag-CD133, using equal cell numbers as input for vesicle preparation., cell viability was determined after 48 hours of incubation. (G) PLC cells were treated with CD133⁺ vesicle isolated from MEKi treated PLC cells. IgG vesicles were used to exclude the non-specific binding. Flowthrough is the un-bound vesicles. CD133+ vesicles were purified by the combination of fractionation and immunoprecipitation. Statistical significance is determined by Student’s t-test. Data were represented as mean ± SEM. *p<0.05, **p<0.01, ***p<0.001, ns: no significance.

To rule out the effects of other vesicle types, we compared vesicles from control cells and CD133-overexpressing cells. Vesicles derived from CD133-overexpressing cells significantly enhanced cell viability compared to those from control cells (Figure 2F), further confirming the role of CD133^+^ vesicles in promoting cell proliferation. Moreover, vesicles isolated from density fractionation followed by immunoprecipitation reveled that treatment with CD133^+^ vesicles significantly increased cell proliferation compared to that of IgG control, whereas vesicles in the flowthrough exhibited the opposite effect (Figure 2G). Taken together, CD133^+^ vesicles promote cell proliferation *in vitro*.

### MEKi-induced CD133 upregulation is mediated by YAP activation

Having established that CD133^+^ vesicles are distinct from EVs and their role in promoting cell proliferation, we next explored the regulatory mechanisms underlying CD133^+^ vesicle formation. CD133 induction was observed approximately 24 hours post MEKi treatment at both the protein and mRNA levels (Supplementary Figure 7A-7D). Concurrently, the Hippo-YAP signaling pathway downstream gene, such as *cyr61*, exhibited a similar induction pattern (Supplementary Figure 7E-7F). This observation prompted us to examine whether MEKi treatment activates YAP, thereby contributing to CD133 induction. To confirm YAP signaling activation in response to MEKi, we measured YAP phosphorylation (pYAP) levels over the course of treatment and observed a significant decrease in the ratio of pYAP to total YAP, indicating YAP activation (Figure 3A-3D). Immunostaining revealed a marked increase in nuclear YAP localization following MEKi treatment compared to the control (Figure 3E-3F’’’’), further confirming YAP activation in response to MEKi.

**Figure 3.**
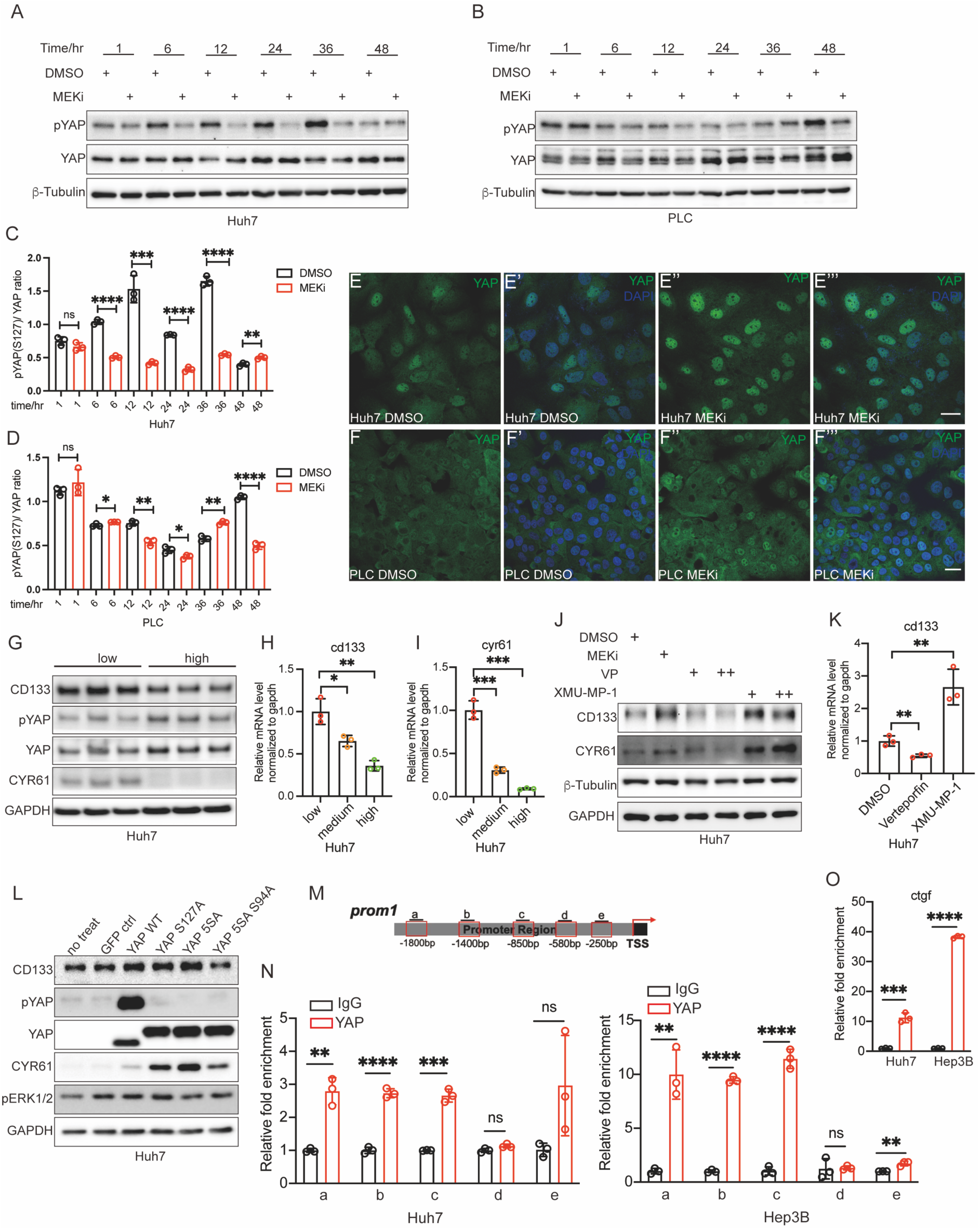
MEK inhibition activates YAP to directly induce CD133 expression. (A) Protein levels of pYAP and YAP were analyzed by western blotting in Huh7 cells treated by MEKi at different time points. (B) Protein levels of pYAP and YAP were analyzed by western blotting in PLC cells treated by MEKi at different time points. (C) The ratio of phosphorylated YAP (pYAP) to total YAP signal intensity was quantified in Huh7 cells treated with MEK inhibitor at various time points. (D) The ratio of phosphorylated YAP (pYAP) to total YAP signal intensity was quantified in PLC cells treated with MEK inhibitor at various time points. (E-E’’’) Immunofluorescence analysis of YAP localization in Huh7 cells. Huh7 cells were treated with (E- E’) DMSO or (E’’-E’’’) MEKi for 24 hours followed by immunostaining for YAP (green), DAPI (blue). MEK inhibition promotes nuclear translocation of YAP. Scale bar 25 μm. (F-F’’’) Immunofluorescence analysis of YAP localization in PLC cells. PLC cells were treated with (F-F’) DMSO or (F’’-F’’’) MEKi for 24 hours followed by immunostaining for YAP (green), DAPI (blue). MEK inhibition promotes nuclear translocation of YAP. Scale bar 25 μm. (G) Protein levels of CD133, pYAP, YAP and CYR61 were analyzed in Huh7 cells at different cell density. CD133 level is decreased in high density condition which consistent with CYR61 level. (H-I) mRNA levels of CD133 (A) and cyr61 (B) were analyzed by qPCR at different cell density in Huh7 cells. (J) Protein levels of CD133 and CYR61 were analyzed when Huh7 cells were treated with MEKi, VP and XMU-MP-1 respectively. VP could inhibit YAP-TEAD binding, which inactivate YAP signaling activity, XMU-MP-1 is a MST1/2 kinases inhibitor that induces YAP activation. (K) mRNA expression level of CD133 was analyzed when treating VP and XMU-MP-1 in Huh7 cells. (L) Protein levels detected by western blot in YAP-WT, YAP-S127A, YAP-5SA and YAP-5SA/S94A overexpression in Huh7 for 48 hours. (M) Diagram of the cd133 promoter region and the detection region for different qPCR primers. (N-O) ChIP-qPCR results showing the binding of YAP to CD133 promoter region in Huh7 and Hep3B cell. Protein bound chromatin was immunoprecipitated with YAP antibody, and IgG was used as a negative control (N). ctgf, a well-known target gene of the YAP-TEAD transcriptional complex, was used as a positive control (O). Statistical significance is determined by Student’s t-test. Data were represented as mean ± SEM. *p<0.05, **p<0.01, ***p<0.001, ns: no significance.

To elucidate the regulation of CD133 by YAP, we examined the effect of cell density on CD133 levels, since low cell density promotes YAP nucleus translocation and activation of downstream targets, whereas high cell density suppresses YAP activity (25, 27, 29, 31). We observed that high cell density markedly reduced CD133 expression across multiple cell lines (Figure 3G-3I, Supplementary Figure 8A-8C), consistent with the downregulation of the classical YAP downstream target, *cyr61*. Additionally, serum stimulation in MCF10A cells induced CD133 expression alongside *cyr61*(25)(Supplementary Figure 8F-8G), indicating that YAP activation positively regulates CD133.

To directly assess the role of the Hippo-YAP signaling pathway in regulating CD133 expression, we treated cells with verteporfin (VP), an inhibitor of YAP-TEAD interaction that blocks YAP activation, and with XMU-MP-1, an MST1/2 kinases inhibitor that induces YAP activation (25, 26, 31, 35). Treatment with VP significantly decreased CD133 levels, while XMU-MP-1 enhanced CD133 expression, further supporting that YAP activity regulates CD133 levels (Figure 3J-3K, Supplementary Figure 8D-8E). Furthermore, transient transfection with constitutively activated YAP (YAP-5SA) significantly increased CD133 expression, while transfection with the TEAD-binding-deficient YAP mutant (YAP-5SA/S94A) resulted in decreased CD133 levels. These findings indicate that activated YAP is a key regulator of CD133 expression, and this regulation depends on YAP-TEAD binding (Figure 3L, Supplementary Figure 8H-8M).

Moreover, we conducted chromatin immunoprecipitation (ChIP)-qPCR assays in both Huh7 and Hep3B cell lines to determine whether the YAP-TEAD complex directly binds to the CD133 promoter region (Figure 3M-3O). Several potential YAP–TEAD binding motifs were identified in the CD133 promoter region (Figure 3M). ChIP-qPCR results confirmed that the YAP-TEAD complex binds to multiple sites within the CD133 promoter region in both cell lines (Figure 3N-3O). Collectively, these findings demonstrated that MEKi induces YAP activation, and activated YAP, in turn, directly targets and upregulates CD133.

### YAP knockdown decreases CD133 expression and vesicular CD133, potentiating MEKi sensitivity

To clarify the role of YAP in regulating CD133, we performed YAP knockdown experiments using short hairpin RNA (shRNA). Silencing YAP reduced basal CD133 expression and abolished MEKi-induced CD133 upregulation (Figure 4A, 4B). Functionally, the combination of YAP knockdown and MEKi treatment significantly inhibited cancer cell proliferation in the PLC stable cell line compared to MEKi treatment alone (Figure 4C). Similarly, CD133 knockdown combined with MEKi treatment produced a similar inhibitory effect on cell proliferation compared to MEKi treatment alone (Figure 4D). Taken together, YAP mediated regulation of CD133 contributes to MEKi resistance and may offer a therapeutic target in liver cancer.

**Figure 4.**
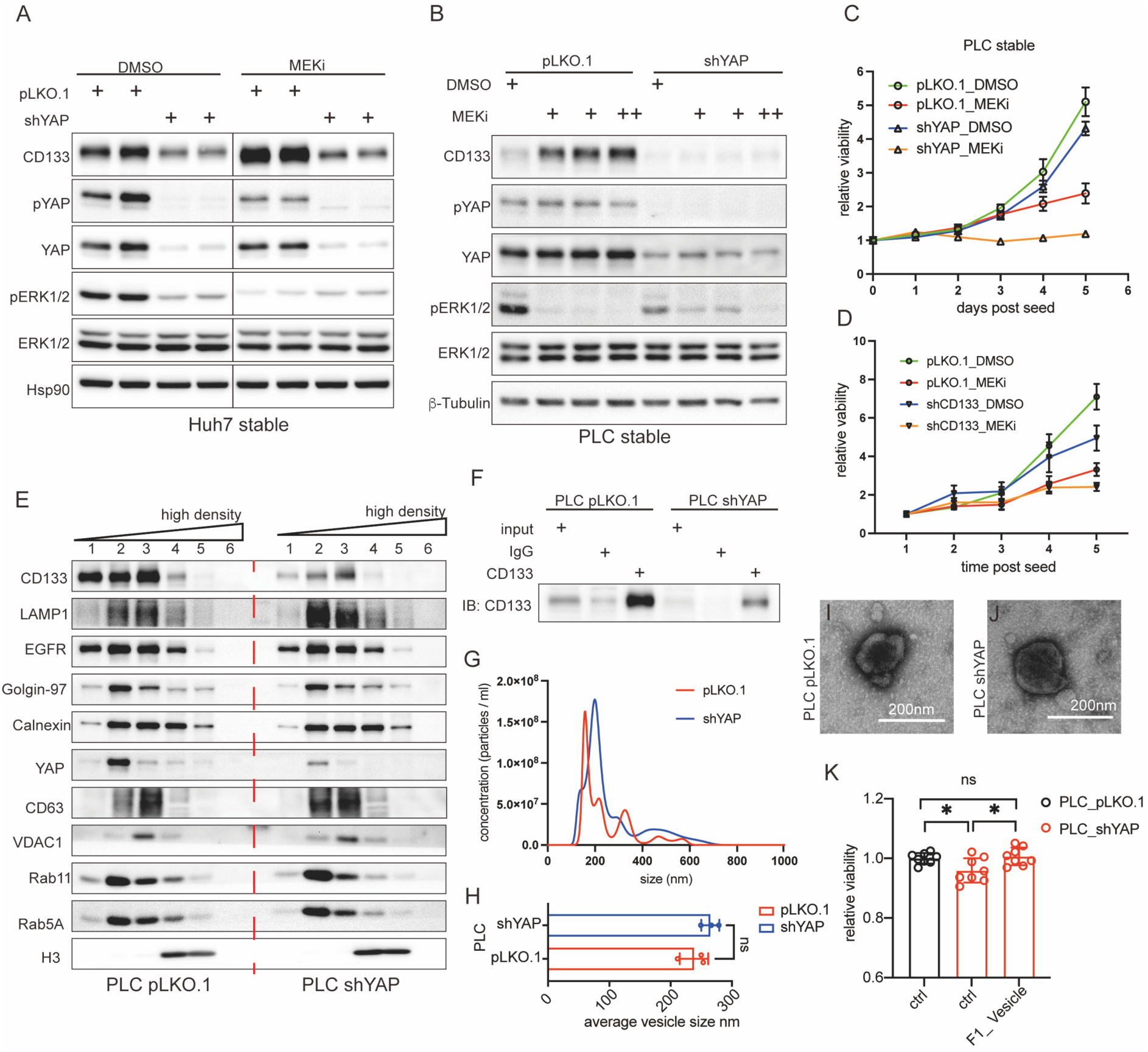
YAP knockdown decreases CD133 expression and vesicular CD133, potentiating MEKi sensitivity. (A) Protein levels of CD133, pERK1/2, ERK1/2, pYAP and YAP were analyzed by western blotting in Huh7 pLKO.1 and Huh7 shYAP stable cell lines in the background of DMSO and MEKi treatment. (B) Protein levels of CD133, pERK1/2, ERK1/2, pYAP and YAP were analyzed by western blotting in PLC pLKO.1 and PLC shYAP stable cell lines in the background of DMSO and MEKi treatment. (C) Cell proliferation assay detected by cck8 when treating the PLC pLKO.1 and PLC shYAP stable cell lines with DMSO or MEKi. Five biological repeats per group. (D) Cell proliferation assay detected by cck8 when treating the PLC pLKO.1 and PLC shCD133 stable cell lines with DMSO or MEKi. Five biological repeats per group. (E) Cell fractions from PLC pLKO.1 control and shYAP stable cells under the MEKi treatment were analyzed by western blotting. Fraction 1 to 6 goes from low density to high density. Same cell number were used in two conditions. CD133 is decreased in the top fractions in shYAP stable cells. EGFR is the plasma membrane marker, Calnexin is an ER marker, Golgin-97 is a Golgi marker. (F) Vesicle IP of CD133^+^ vesicle from PLC pLKO.1 and PLC shYAP stable cells under the MEKi treatment condition. Same cell numbers were used in both cell lines. (G) NTA of indicated vesicles of the top fraction from PLC pLKO.1 and PLC shYAP stable cells showing size distribution (n=3 biological replicates). (H) Statistics of the average vesicle size from NTA measurement in the top fraction from both PLC pLKO.1 and PLC shYAP stable cells under the MEKi treatment. Statistical significance is determined by Student’s t- test. Data were represented as mean ± SEM. ns: no significance. (I-J) Representative electron microscope image of the vesicles from top fraction of both PLC pLKO.1 and PLC shYAP stable cells. Scale bar, 200nm. (K) Cell viability of PLC_shYAP stable cells were determined with or without CD133^+^ vesicles enriched fractions addition after 36 hours treatment. Cell viability was normalized with the PLC_pLKO.1 stable cells without vesicle addition. Statistical significance is determined by Student’s t-test. Data were represented as mean ± SEM. *p<0.05, ns: no significance.

Based on the finding that CD133 protein is regulated by YAP, we hypothesized that YAP knockdown may also affect CD133^+^ vesicles. To test this, we isolated vesicles from mock and shYAP stable cell lines to isolate vesicles. Fractionation analysis revealed a significant reduction of CD133 in the top fraction following YAP knockdown, indicating decreased vesicular CD133 levels (Figure 4E). This reduction was further validated by vesicle immunoprecipitation (Figure 4F). Importantly, measurements of average vesicle sizes and vesicle size distribution in the top fraction revealed no significant changes between mock and shYAP (Figure 4G-4H), nor were there any changes in vesicle morphology (Figure 4I, 4J). These findings demonstrated that YAP knockdown reduced vesicular CD133 levels without altering vesicle size or morphology. Furthermore, we examined the effect of CD133^+^ vesicles on shYAP stable cell lines. While YAP knockdown reduced cell proliferation, treatment with CD133^+^ vesicles enriched fraction restored cell viability of shYAP cells to levels comparable to control levels (Figure 4H). This result indicates that CD133 functions downstream of YAP and that CD133^+^ vesicles are sufficient to rescue cell proliferation despite YAP suppression, highlighting their critical role in sustaining cell growth.

### CD133 is upregulated in YAP-5SA induced liver cancer *in vivo*

To further explore the regulatory relationship between YAP and CD133 *in vivo*, we employed hydrodynamic tail vein injection (HTVi) to deliver constitutively activated YAP (YAP-5SA) into the liver to induce liver tumor formation (Figure 5A) (25, 35, 36). Liver tissues were collected four months post injection. While livers from GFP control mice appeared normal, tumor nodules were clearly developed in the YAP-5SA group. The liver-to-body weight ratio was increased in YAP-5SA mice compared to controls (Figure 5B). In addition, both tumor size and tumor number were markedly increased (Figure 5C, 5D). Analysis of these tissue samples revealed elevated CD133 expression at both the mRNA and protein levels (Figure 5E, 5I). Additionally, canonical YAP downstream targets *cyr61*, *ccnd1*, *sox9* and *survivin* (25, 26, 28, 29) were all robustly induced in the tumor regions (Figure 5F-5I).

**Figure 5.**
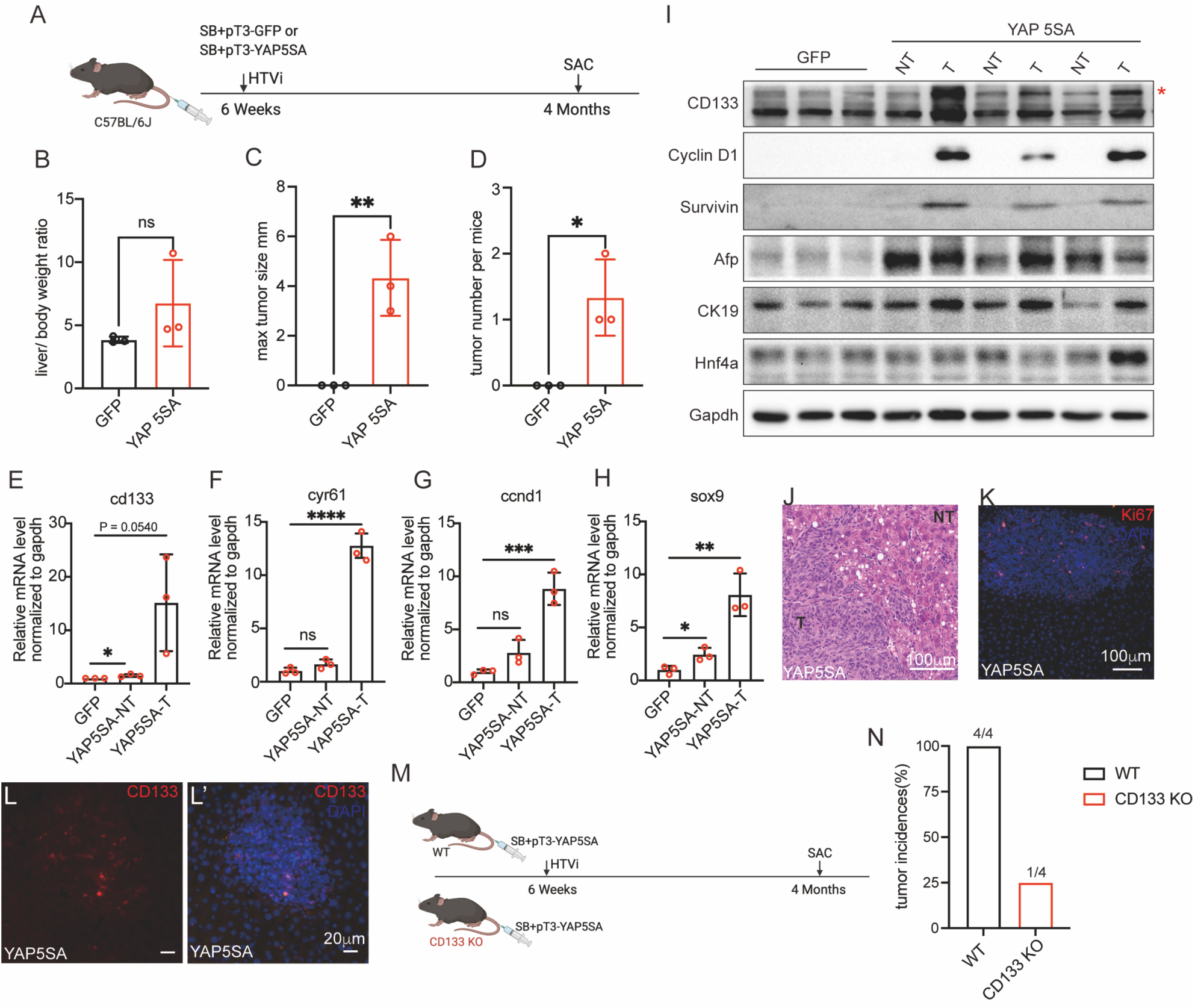
CD133 is upregulated in YAP-5SA induced liver cancer *in vivo*. (A) Schematic of liver tumorigenesis model by the hydrodynamic tail vein injection. Six-week-old WT mice received SB along with pT3-GFP or pT3-YAP-5SA plasmids by hydrodynamic tail vein injection (HTVi). Livers were harvested after hydrodynamic transfection for 4 months to determine tumor burden. (B-D) Tumor burdens were calculated by (B) liver/body weight ratio, (C) maximal tumor sizes, or (D) numbers of tumor nodules. Data were represented as mean ± SEM. **p<0.01, *p<0.05, ns: no significance. n=3. (E-H) mRNA level of CD133, cyr61, ccnd1 and sox9 were analyzed by qPCR from liver tissue. Cyr61, ccnd1 and sox9 are classic Hippo-YAP signaling downstream targets. Data were represented as mean ± SEM. **p<0.01, *p<0.05, ns: no significance. n=3. (I) Protein expression levels analyzed by western blotting in liver tissues from GFP control mice, YAP-5SA mice. CD133 is significantly upregulated in the tumor tissue regions compared with the non-tumor tissue and GFP controls. * indicates the specific CD133 bands. (J) Representative hematoxylin and eosin staining of liver sections from YAP-5SA liver tissue. Scale bar, 100μm. (K) Representative Ki67 staining of liver sections from YAP-5SA liver. Ki67, red. DAPI, blue. Ki67 signal is much higher in tumor regions compared with the non-tumor region. Scale bar, 100μm. (L) Representative CD133 staining of liver sections from YAP-5SA liver. CD133, red. DAPI, blue. CD133 signal is much higher in tumor regions compared with the non-tumor region. Scale bar, 20μm. (M) Schematic of liver tumorigenesis model by the hydrodynamic tail vein injection. SB along with pT3- YAP-5SA plasmids were injected into WT mice or CD133KO mice by tail vein injection. Livers were harvested after hydrodynamic transfection for 4 months to determine tumor burden. (N) Tumor incidences, as percentages of total mice (N = 4), with the number of tumor- bearing mice over the total number of mice with indicated genetic background as (M) post HTVi.

**Figure 6.**
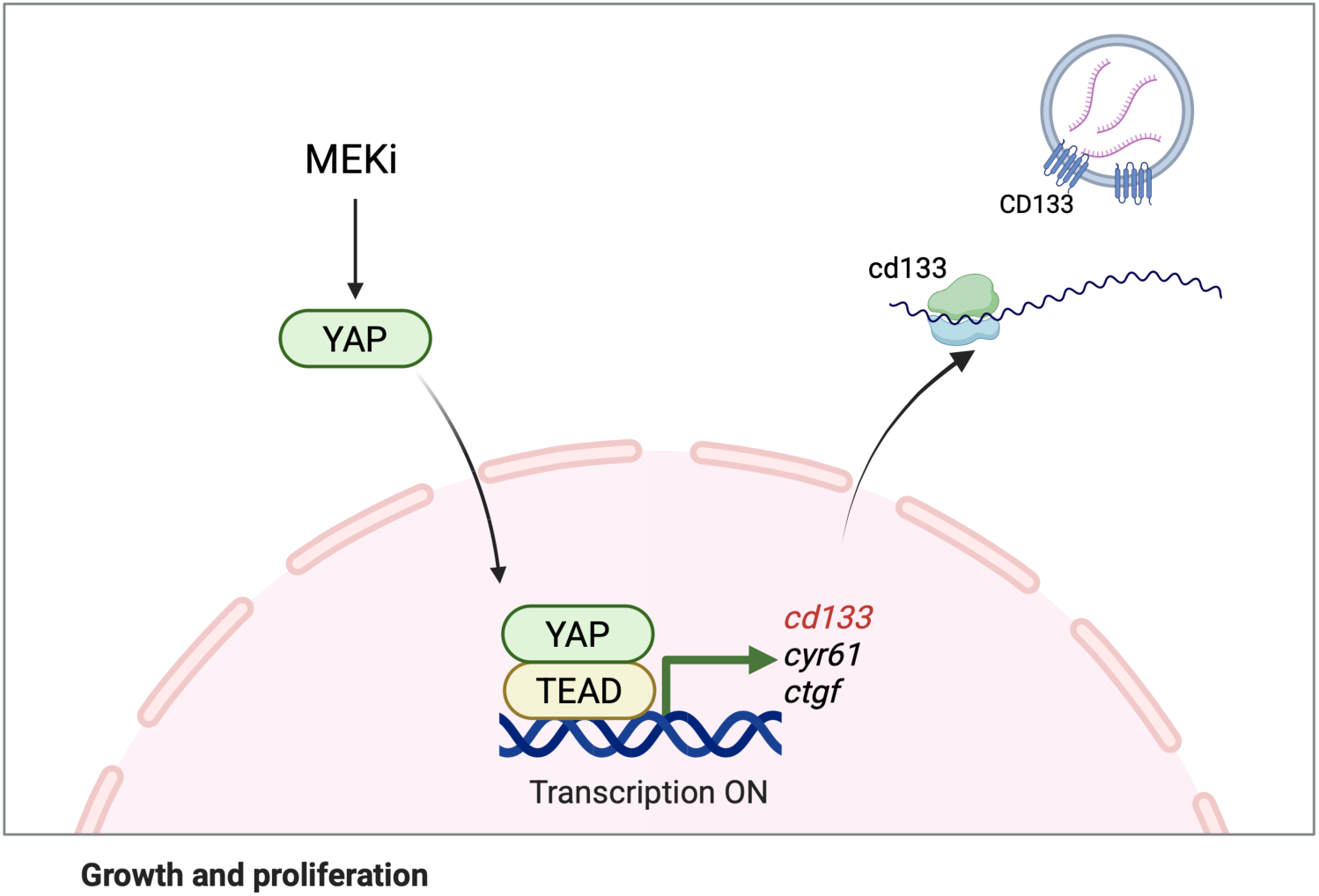
A model for MEKi induced CD133^+^ vesicle production. Treatment with MEKi impairs cancer cell proliferation. In response, cancer cells activate the Hippo-YAP signaling pathway, resulting in nuclear translocation of YAP and subsequent transcriptional upregulation of CD133. Elevated CD133 expression promotes the formation of CD133⁺ vesicles, which contribute to adaptive resistance by mitigating MEKi-induced proliferative defects. This compensatory mechanism supports the continued proliferation of cancer cells under MEKi treatment.

Histology staining confirmed tumor presence (Figure 5J), and immunofluorescence analysis demonstrated significantly increase in Ki67 levels in tumor regions of YAP-5SA mice livers compared to non-tumor areas (Figure 5K). Notably, CD133 expression was elevated in tumor regions relative to the surrounding non-tumor areas in liver of YAP-5SA mice (Figure 5L-5L’). Additionally, a parallel HTVi experiment performed in CD133 knockout (KO) mice resulted in reduced tumor formation (Figure 5M, 5N), unlike the CD133 wild type controls. These results demonstrated that CD133 is critical for YAP-5SA-driven liver cancer formation *in vivo*.

## Discussion

The RAS-ERK signaling pathway plays an indispensable role in regulating cell proliferation and cancer progression (4, 5). SHP2 is required for the fully activation of RAS-ERK signaling (5). A previous study has identified a novel CD133⁺ intercellsome structure emerging in the SHP2 knockout liver following partial hepatectomy; however, the properties and underlying mechanisms of this phenomenon remain largely unexplored (7). In this study, we characterized the properties, function, and regulatory mechanisms of this novel population of intracellular CD133^+^ vesicles, which were strongly induced in response to proliferation defects caused by RAS-ERK pathway dysregulation. Here, we provided the first comprehensive isolation and molecular characterization of CD133⁺ vesicles. Our findings revealed that Hippo-YAP signaling pathways governs their formation and function, offering new insights into how cells respond to proliferative stress via the RAS-ERK pathway (Supplementary Figure 9).

We demonstrated that CD133 expression is significantly upregulated by MEKi treatment across multiple cell lines, leading to increased production of CD133^+^ vesicles. Although these vesicles share a similar size profile to conventional EVs, their RNA cargo, including both mRNAs and microRNAs, is distinct. This molecular difference indicates that CD133⁺ vesicles are not typical EV precursors but represent a novel class of intracellular vesicles with unique molecular characteristics and biological roles. While preliminary RNA profiling supports this distinction, a more comprehensive analysis such as comprehensive proteomic profiling and lipid profiling are required to fully understand the molecular functions and signaling capacities of these vesicles. In addition, it’s essential to identify additional unique molecular markers specific to these vesicles to further characterize their biological properties and functional roles.

Further investigation into the regulatory mechanisms underlying CD133^+^ vesicle production upon MEKi exposure revealed that MEKi could activate the Hippo-YAP signaling pathway. Specifically, activated YAP directly upregulated CD133 expression, thereby promoting the formation of CD133^+^ vesicles in response to impaired proliferative signals. These findings not only enhance our understanding of the biogenesis regulatory mechanism of CD133^+^ vesicles but also provide insights into potential mechanisms of drug resistance in liver cancer treatment. The activation of the Hippo-YAP pathway and subsequent upregulation of CD133^+^ vesicle production may represent an adaptive response that enables cancer cells to survive proliferative stress, thereby contributing to therapeutic resistance. YAP-5SA induced liver cancer model confirmed that YAP activation could induce CD133 *in vivo*. However, further study should be conducted *in vivo* to study whether upregulated CD133 formed vesicles and how these vesicles communicated between cancer cells. Understanding the role of CD133⁺ vesicles in liver cancer progression and therapy resistance could provide novel therapeutic targets for overcoming drug resistance in liver cancer.

Isolating pure CD133⁺ vesicles presents a significant technical challenge due to their intracellular origin and potential co-isolation with other vesicle populations. To address this, we have developed a novel protocol that integrates high-resolution density gradient fractionation with immunoprecipitation. This combined approach allows us to effectively isolate CD133^+^ vesicles, enhancing the accuracy and reproducibility of downstream analyses. Nonetheless, some non-specific enrichment may still occur, warranting further optimization. Continued efforts to develop more convenient and efficient isolation methods are crucial, as such advances would greatly enhance the specificity, scalability, and reproducibility of CD133^+^ vesicle isolation, thereby facilitating downstream application and mechanism study.

Beyond defining their composition and regulation, an important aspect is to understand how CD133⁺ vesicles were transported in intercellular communication. We are currently exploring how these vesicles are transferred between cells, a process that could underlie broader signaling networks in response to proliferative stress. Gaining mechanistic insight into this trafficking process will be critical to uncovering the biological relevance of CD133⁺ vesicles in both normal physiology and disease contexts.

Our study provides significant insight into the biogenesis and regulation of CD133⁺ vesicles in response to MEKi treatment. We identified YAP activation as a critical upstream driver of CD133⁺ vesicle production, highlighting a mechanistic link between proliferative stress and vesicle formation. These findings suggest that modulating the YAP-CD133⁺ vesicle axis may represent a promising approach for combination therapies aimed at limiting tumor adaptation and recurrence. Ongoing efforts to explore the downstream impact of these vesicles will further expand our understanding of their role in cancer biology.

In summary, our findings represent a critical step toward understanding the biological functions and regulatory mechanisms of CD133^+^ vesicles. By characterizing their unique molecular composition and identifying YAP-driven induction under proliferative stress, we establish CD133⁺ vesicles as a distinct intracellular vesicle population. Continued investigation into their roles in intercellular communication and therapeutic adaptation, particularly in liver cancer, holds promise for uncovering new strategies to overcome drug resistance.

## Methods

### Cell culture

PLC/PRF/5, Huh7, Hep3B, HepG2, MDA-MB-231, CaCo2, MiaPaca2, Capan2, HeLa cells were cultured in DMEM medium supplemented with 10% fetal bovine serum. The MCF10A cells were cultured in DMEM-F12 medium (Gibco) with 5% horse serum (Gibco) and 1% penicillin-streptomycin supplemented with 20 ng/mL human EGF (Gemini), 0.5 μg/mL hydrocortisone, 100 ng/mL cholera toxin (Sigma-Aldrich), and 10 μg/mL human insulin (Sigma). BxPc3 cells were cultured in RPMI-1640 (Gibco) with 10% fetal bovine serum. Panc10.05 cells were cultured in RPMI-1640 (Gibco) with 10% fetal bovine serum and 10 μg/mL human insulin (Sigma). All cell cultures are supplied with 1% Penicillin-Streptomycin (Gibco) and cultured in 37°C humidified environment containing 5% CO_2_ environment. Inhibitor treatments followed previously reported protocols(7). Briefly, cells were treated with SHP2i (SHP099, chemietek, 5μM), MEKi (trametinib, APExBIO, 10nM), Refametinib (MCE, 1μM), Ulixertinib (MCE, 1μM), Selumetinib (MCE, 1μM), Vertepofin (MCE, 1μM), XMU-MP-1 (MCE, 1μM) according to the specified concentrations (25). Cells were treated for 48hrs or 72hrs. All cell lines have been tested for mycoplasma contamination.

### Animals

All mice were maintained under a 12-hour light/dark cycle with free access to water and mouse chow food. All animal protocols were approved by the institutional animal care and use committee of the University of California, San Diego. Mouse liver tumors were induced by hydrodynamic tail vein injection (HTVi) of pT3-YAP-5SA together with the sleeping beauty transposase, as described previously (7, 25, 36). All plasmid DNAs were diluted in phosphate-buffered saline (PBS) and injected at 0.1 mL/g body weight through the tail vein within 5-7 seconds. CD133 knockout mice (B6N;129S-Prom1^tm1(cre/ERT2)Gilb/^J) were purchased from the Jackson Laboratory and breed back with C57BL/6 wild type mice to C57BL/6 genetic background (7).

### Immunoblotting

Proteins were extracted in RIPA buffer, and immunoblotting was performed using standard protocols as previously described., specific primary antibodies used in the experiments are listed here. mouse CD133 (14133182, eBioscience), human CD133 (86781, CST), human CD133 (130092395, miltenyi), CD133 (18470, proteintech), CD133 (66666, proteintech), β-Tubulin (66240, proteintech), pERK1/2 (4370, CST), ERK1/2 (4695, CST), pYAP (13008, CST), YAP (14074, CST), CD63 (GTX41875, GeneTex), Rab11 (5589, CST), Calnexin (2679, CST), VDAC1 (14734, abcam), H3 (4620, CST), Rab5A (46449, CST), EGFR (4267, CST), LAMP1 (21997, proteintech), Golgin-97 (13192, CST), Cyr61 (PA5-78022, Invitrogen) and Golgi58 (ab27043, abcam). Images are visualized using the ECL system (34580, Thermo Fisher) followed by Bio-Rad ChemiDoc imaging system. Intensity of the bands in the image was measured using ImageJ software (NIH).

### RT-qPCR

RNAs extracted by Trizol following standard protocols were reverse transcribed into cDNA using Hiscript RT supermix kit (R323, Vazyme) and quantitative PCR was performed using Taq Pro Universal SYBR qPCR Master Mix (Q712, Vazyme) on Bio-Rad CFX Duet Real-Time PCR system. Primers used in this study are listed here:

human CD133 (Forward: TCGAATGGATTCGGAGGACG; Reverse: TGTTGTGATGGGCTTGTCAT),

human gapdh (Forward: GAGTCAACGGATTTGGTCGT; Reverse: GACAAGCTTCCCGTTCTCAG),

human cyr61 (Forward: AGCCTCGCATCCTATACAACC; Reverse: TTCTTTCACAAGGCGGCACTC),

mouse gapdh (Forward: CTCCCACTCTTCCACCTTCG; Reverse: TAGGGCCTCTCTTGCTCAGT),

mouse cyr61 (Forward: AGAGGCTTCCTGTCTTTGGC; Reverse: CTCGTGTGGAGATGCCAGTT),

mouse CD133 (Forward: TGTTGGTGCAAATGTGGAAAAG; Reverse: ATTGCCATTGTTCCTTGAGCAG).

### Isolation of CD133^+^ vesicles

To isolate CD133^+^ vesicles, cultured cells were treated with a MEK inhibitor (MEKi) for three days to promote vesicle production (7). Following treatment, the culture medium was discarded, and the cells were washed three times with cold PBS to remove residual medium. The cells were then scraped off the culture dish and resuspended in an appropriate volume of ice-cold Dulbecco’s PBS (DPBS, 14190144, Thermo Fisher). To lyse the cells, the suspension was processed on ice using a Dounce homogenizer, and proteinase inhibitor (1:500, 78441, Thermo Fisher) was added to prevent proteolytic degradation. The lysis process was monitored microscopically, and once most nuclei were released, the lysate was subjected to differential centrifugation to remove cellular debris and organelles. The lysate was centrifuged at *300 × g*, *2000 × g*, and *12,000 × g* at 4°C to pellet nuclei, mitochondria, and large organelles, respectively, and the supernatant was retained for further analysis. To purify vesicles, a discontinuous iodixanol (OptiPrep, D1556, Sigma-Aldrich) gradient (12%–36%) was prepared using ice-cold DPBS, and the lysate supernatant was mixed with iodixanol to achieve a final concentration of 12%. This mixture was loaded on top of the pre-formed gradient and ultracentrifuged at *120,000 × g* for 15 hours at 4°C using the SW41 Ti Swinging Bucket rotor (k factor of 124, Beckman Coulter). After centrifugation, six individual fractions (1.6 mL each) were carefully collected from the top of the gradient. The top fraction, enriched with CD133^+^ vesicles, was subjected to immunoprecipitation using a CD133-PE antibody (12133842; Invitrogen), followed by further purification with the Anti-PE multi-Sort Kit (130090757, Miltenyi) to isolate pure CD133^+^ vesicles. For Nanoparticle Tracking Analysis (NTA), vesicles were diluted in particle-free PBS. For immunoblotting, vesicles or fractions were lysed in RIPA buffer for 10 minutes on ice (25). For RNA extraction, vesicles were processed using RNeasy Micro Kit (74004, QiaGen) following the standard protocol.

### Electron Microscopy

Negative staining. Freshly prepared fractions or pure vesicles were loaded onto pre-discharged grids (0.1754-F, Electron Microscope) and subsequently washed three times with double-distilled water for 30 seconds each. The samples on the grid were then transferred to a 2% uranyl acetate (UA) solution for 3 seconds, followed by incubation with another drop of UA for 1 minute. After staining, the grid was air-dried on tweezers for approximately 20 minutes before further analysis.

Immune-gold labeling staining. Freshly prepared fractions or pure vesicles were first loaded onto pre-discharged grids and blocked with 1% bovine serum albumin (BSA) in PBS for 30 minutes at room temperature. Following blocking, the grids (carbon side down) were incubated in the primary antibody solution for 1 hour at room temperature. Subsequently, the sample grids were washed five times for 30 seconds each. Secondary antibodies conjugated with gold particles of varying sizes were diluted at 1:100 in PBS containing 0.1% BSA and incubated with the grids for 1 hour at room temperature. Finally, similar to the negative staining process, the grids were stained with UA. Images were taken by Tecnai G2 Spirit BioTWIN Electron Microscope at Electron Microscope Core at UCSD.

### Immunofluorescence staining

Cells were cultured in the cover glass in 24 well plates after treating for MEKi for 22 hours. Cells were fixed by 4% PFA for 10 minutes at room temperature. Antibody for YAP (1:500, WH0010413M1, Sigma) was used. All the other steps are following standard procedure as previously described(37, 38). Images were taken with Leica Stellaris 5 Confocal at UCSD School of Medicine Microscopy Core. Immunostaining was performed on fresh frozen liver tissue sections following standard procedure as previously described(7, 31). Liver tissue sections were fixed in 4% PFA overnight at 4 degrees, and antibody for mouse CD133 (1:500, 14133182, eBioscience) was used. All the other steps are following standard procedure as previously described(7, 25, 31).

### Plasmids construction

Human Flag-CD133 and CD133-Flag were constructed into pCDH vector. YAP-5SA was sub-cloned into pT3 vector for hydrodynamic tail vein injection experiment. pcDNA-YAP WT, pcDNA-YAP S127A, pcDNA-YAP 5SA and pcDNA-YAP 5SA/S94A are kind gifts from Dr Xue Feng (Department of Medicine, UCSD, La Jolla, CA). pLKO.1-shYAP (plasmid #27368) was purchased from Addgene (36).

### Cell transfection and viral infection

Plasmid transfection was performed using Lipofectamine 3000 (Invitrogen) following the manufacturer’s instructions. Cells were collected 48 hours or 60 hours post transfection. To generate stable cell lines, lentiviral infection was employed as previously described. Briefly, HEK293T cells were co-transfected with viral vectors and packaging plasmids. After 72 hours, the viral supernatant was harvested, filtered through a 0.45μm filter, and used to infect target cells. Following infection (48 hours later), cells were selected using culture medium containing 4 μg/ml puromycin, which was refreshed every two days.

### Cell Counting Kit-8 (CCK8) assay

Cells were seeded in 96-well plate and the viability of cells was measured by CCK8 assay. For cell growth curve, the CCK8 is measured daily for 5 days. For vesicle treatment assay, cell viability was measured at the first day of treating vesicles and 48hrs post treatment. Briefly, 10 μL of CCK8 solution was added into each well and incubated at 37°C for 4 hours. The absorbance was measured at the wavelengths of 450 nm.

### ChIP-qPCR

ChIP assays were performed as described previously(25). Chromatin was immunoprecipitated with 2 μg antibody of YAP (WH0010413M1, Sigma, kind gift from Dr Xue Feng) or normal mouse IgG (sc-2025, Santa Cruz). DNA immunoprecipitated by the antibodies were detected by RT-qPCR. The primers used were listed below:

CD133-Region a: (Forward: GACCATCTTGCTTCACCCCTT; Reverse: CCTCAGTTTCCTGGTCTGCAA)

CD133-Region b: (Forward: AACTTCTGACCTGCAGGGGC; Reverse: AGCCTCTGAAACCCAGTCTC)

CD133-Region c: (Forward: CCAGTACCTGTTTTATGTTAG; Reverse: ATGGATAGACAGATGAAAGG)

CD133-Region d: (Forward: GGTGAGGACACTTAGACACCG; Reverse: GCAGTGGCTTCACGTTATAG)

CD133-Region e: (Forward: CCGGGATGAGACAGGAGAGT; Reverse: TCCTGTAGAGGTGCCTGCTC)

CTGF: (Forward: TGTGCCAGCTTTTTCAGACG; Reverse:

TGAGCTGAATGGAGTCCTACACA)

### RNA-seq

Total RNA was isolated and purified using RNeasy Micro Kit (74004, QiaGen). Libraries with different indices were multiplexed and loaded on an Illumina NovaSeq6000 instrument according to manufacturer’s instructions. Sequencing was carried out using a 2x150bp paired-end configuration. The reads were mapped to the human reference genome (Ensemble GRCh38.84) using TopHat. The FPKM values were calculated by Cufflinks using default parameters. The quality control, library preparation, and RNA sequencing were completely conducted by IGM core at UCSD.

### Statistical analysis

All of the statistical analyses are one-tailed, unpaired, t-tests with equal variances. All of the experiments are repeated three independent times with similar results unless stated otherwise. Images shown are the representative of images obtained, which are not less than 5. Data were analyzed using the Prism 9 software (GraphPad, San Diego, CA).

## Acknowledgements

The authors thank all members of the Feng laboratory for helpful discussion and technical support throughout the project. The authors thank Dr Wei Ying and Dr Karina Cunha e Rocha (Department of Medicine, UCSD) for the Nanoparticle Tracking Analysis. The authors thank Dr Emily Shizhen Wang and Dr Minghui Cao (Department of Pathology, UCSD) for the Beckman Ultracentrifuge. The authors thank the UCSD School of Medicine Microscopy Core (P30NS047101). The authors thank the UCSD, Cellular and Molecular Medicine Electron Microscopy Core (UCSD-CMM-EM Core, RRID: SCR_022039, S10OD023527) for equipment access and technical assistance. This work was supported by NIH grants (R01DK128320, R01CA236074, R01CA239629 and P01AG073084) to Gen-sheng Feng. Xue Feng was supported by American Diabetes Association Postdoc fellowship (ADA316035- 00001)

## Author contributions

G.F. conceived and supervised this project, analyzed data, and prepared the manuscript. S.Z. performed experiments, analyzed data, and drafted the manuscript; X.F. performed experiments, analyzed data, contribute reagents and drafted the manuscript; E.Q. analyzed the RNA sequencing data; K.K. contributed to reagents and materials.

## Supplementary Figures

**Supplementary Figure 1.**
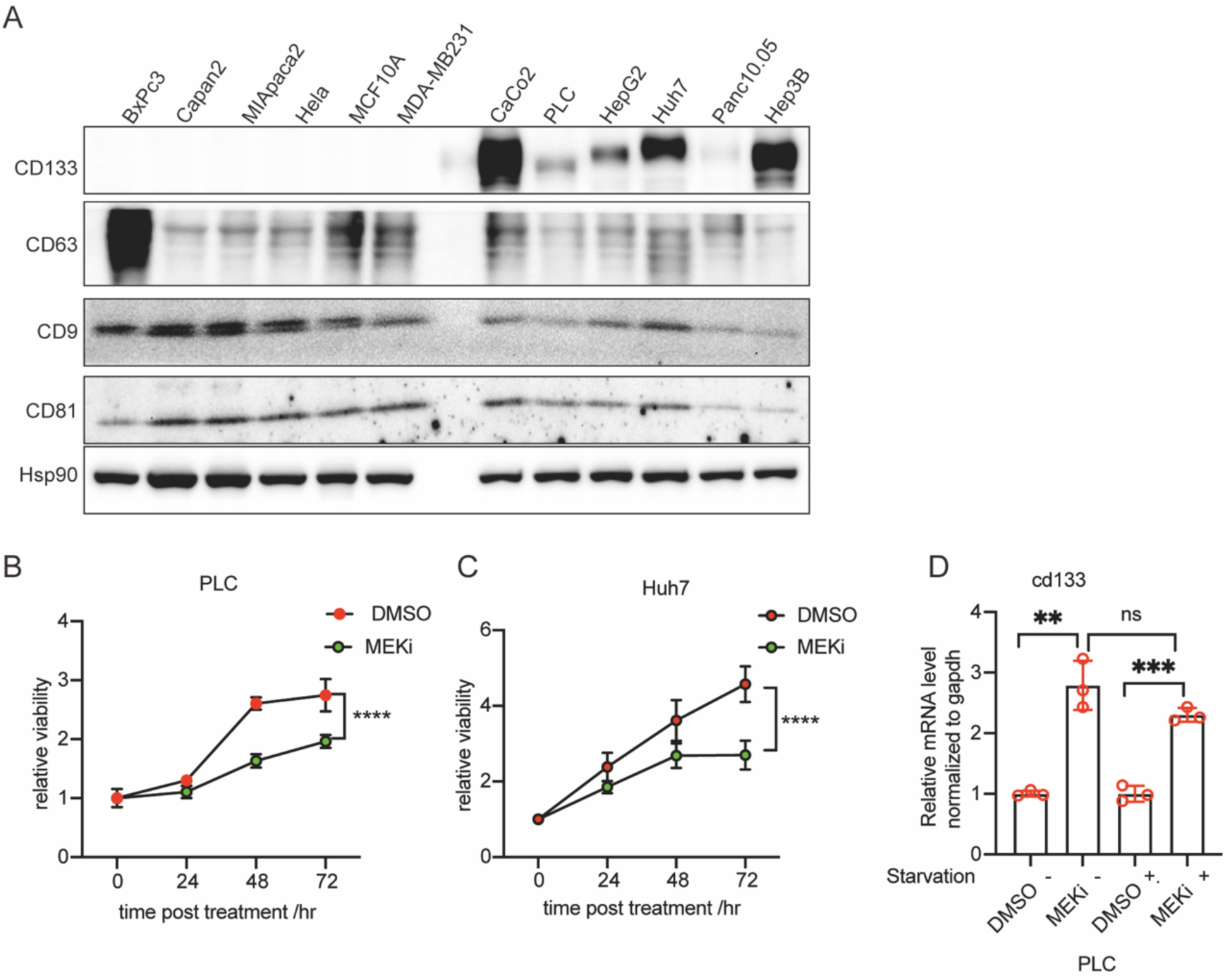
CD133 is highly expressed in liver and colon cancer cells, without specific correlation to the EV marker CD63. (A) Protein levels of CD133, CD63, CD9 and CD81 were analyzed by western blotting in different cancer cell lines. CD63, CD9 and CD81 are common EV markers. (B-C) Cell proliferation Curve of liver cancer cell lines in DMSO control and MEKi treatment. (B) PLC and (C) Huh7 cells were treated with either DMSO (vehicle control) or MEKi. Cell viability was assessed at the indicated time points using the CCK-8 assay. Relative cell viability was normalized to the baseline and plotted over time. MEKi treatment resulted in a marked reduction in cell proliferation compared to the DMSO control in both cell lines. (D) CD133 mRNA level were analyzed by qPCR after MEKi treatment in PLC cells with or without starvation. Statistical significance is determined by Student’s t-test. Data were represented as mean ± SEM. *p<0.05, **p<0.01, ***p<0.001.

**Supplementary Figure 2.**
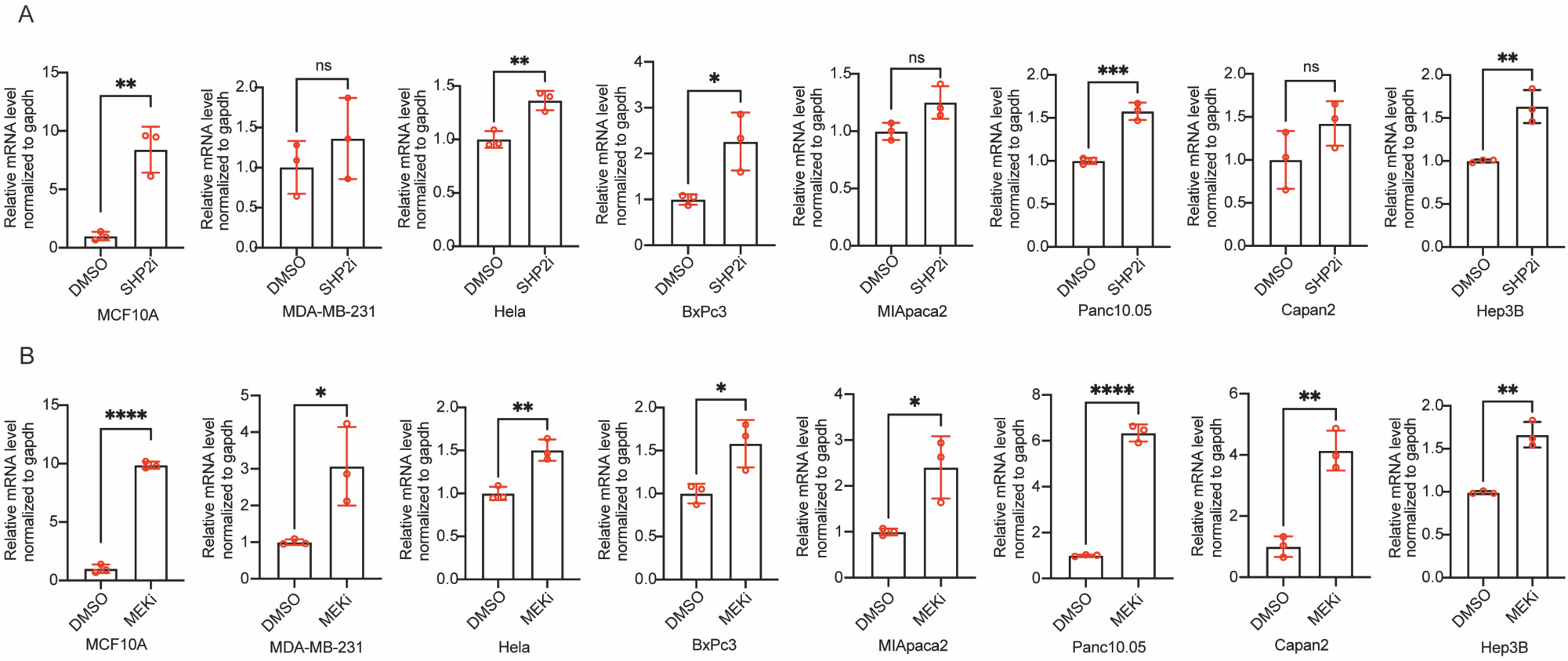
Both SHP2i and MEKi could transcriptionally upregulate CD133 in multiple cancer cell lines. (A) CD133 mRNA level were analyzed by qPCR in multiple cell lines with treated with SHP2i. (B) CD133 mRNA level were analyzed by qPCR in multiple cell lines with treated with MEKi. Statistical significance is determined by Student’s t-test. Data were represented as mean ± SEM. *p<0.05, **p<0.01, ***p<0.001.

**Supplementary Figure 3.**
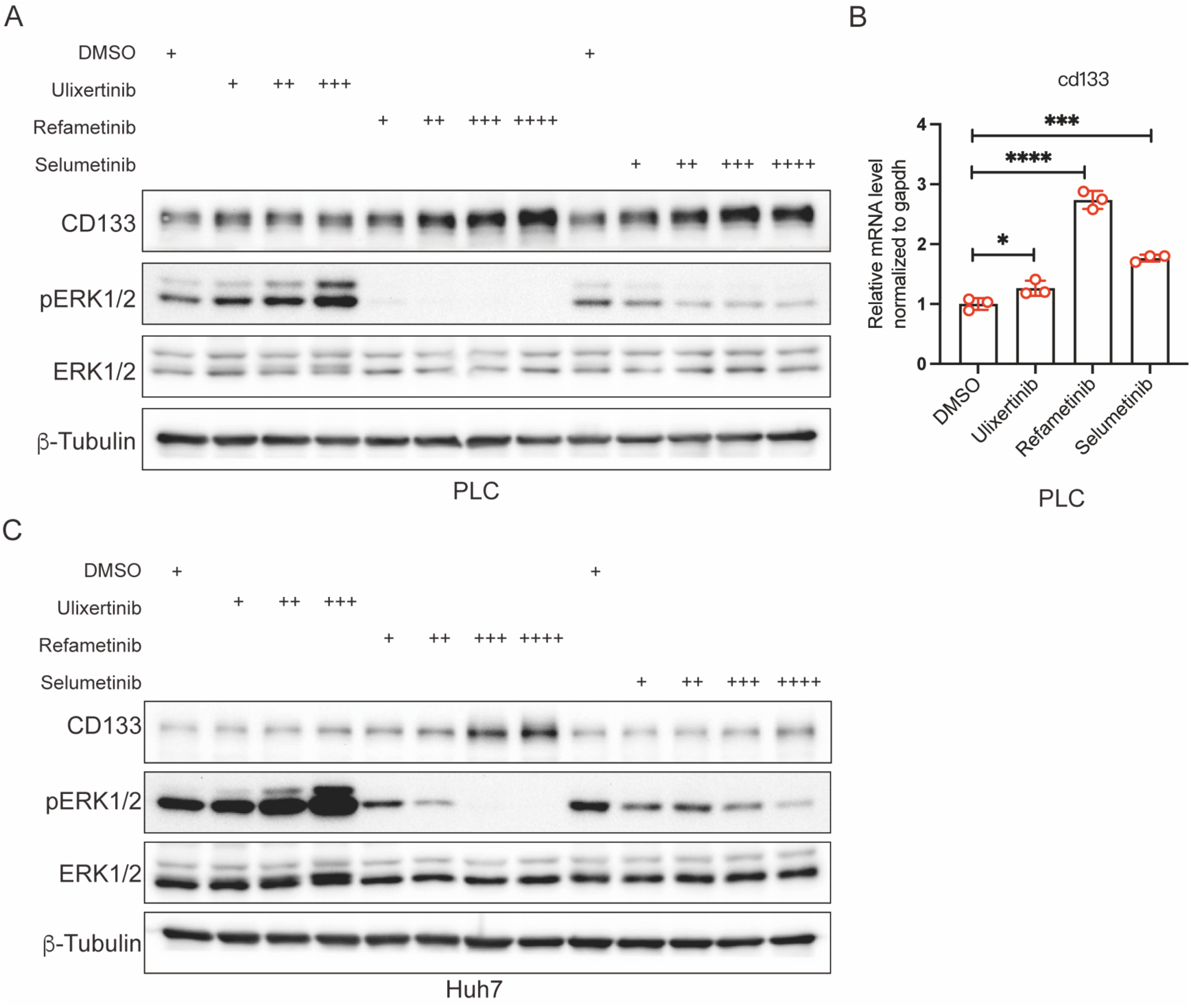
Treatment with various MEK/ERK Inhibitors induces CD133 Expression. (A) Protein levels of CD133 and pERK1/2 were analyzed by western blotting when treating different dosages of MEK/ERK inhibitors in PLC cells. Cells were treated for 48hrs. Ulixertinib is a ERK inhibitor, Refametinib and Selumetinib are MEK inhibitors. (B) CD133 mRNA expression is upregulated in PLC cells following treatment with multiple MEK/ERK inhibitors. PLC cells were treated with inhibitors for 48 hours. Statistical significance is determined by Student’s t-test. Data were represented as mean ± SEM. *p<0.05, **p<0.01, ***p<0.001. (C) Protein levels of CD133 and pERK1/2 were analyzed by western blotting when treating different dosages of MEK/ERK inhibitors in Huh7 cells. Cells were treated for 48hrs. Ulixertinib is a ERK inhibitor, Refametinib and Selumetinib are MEK inhibitors.

**Supplementary Figure 4.**
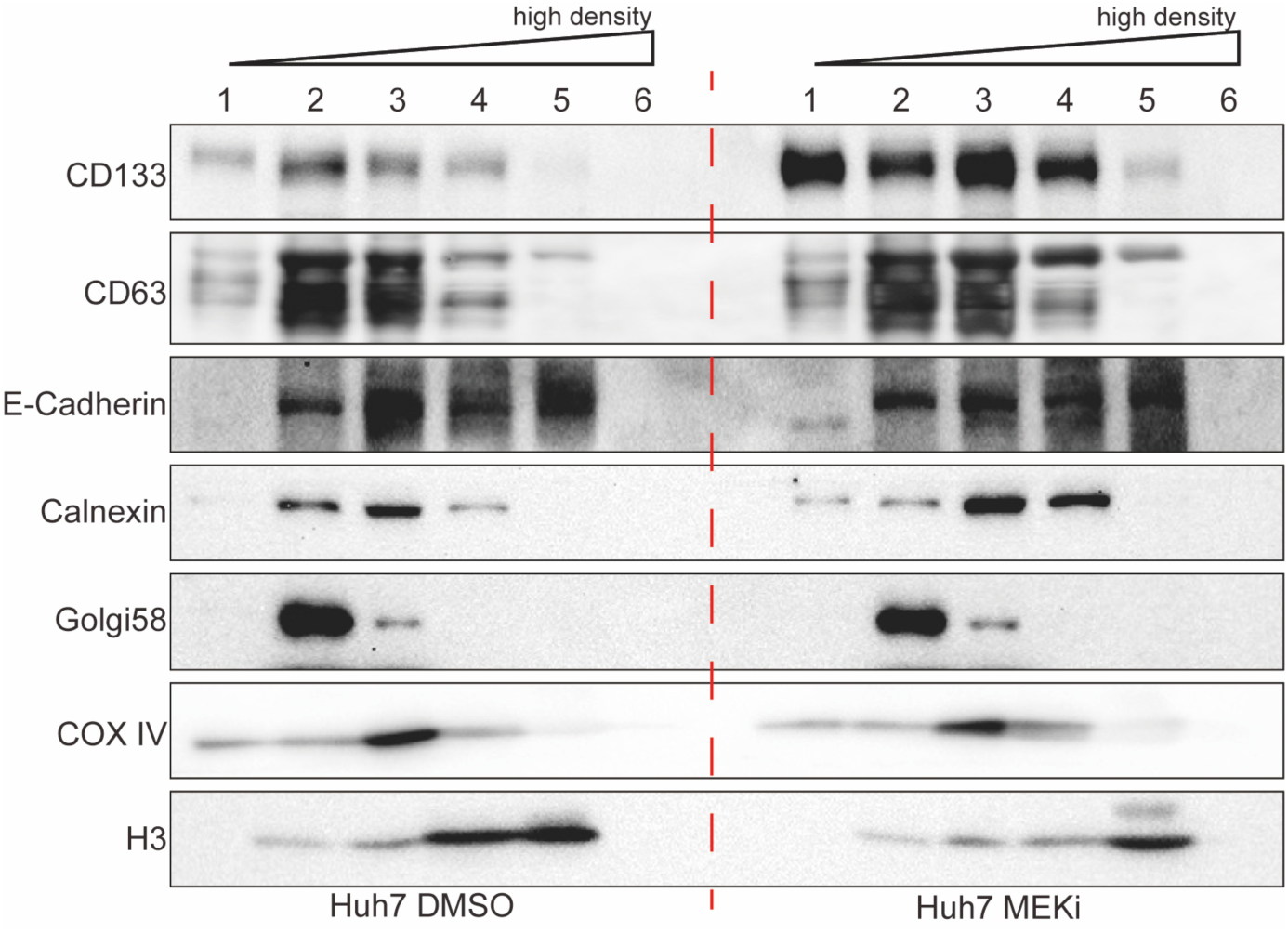
Increased vesicular CD133 in top Fraction of Huh7 Cells. Western blotting showing Huh7 fractionation contents in DMSO and MEKi condition. Fraction 1 to 6 goes from low density to high density. CD63 is a marker for multiple vesicular body, E-Cadherin is a marker for cell membrane, Calnexin is a marker for ER, Golgi58 is a marker for Golgi apparatus, COX IV is a marker for mitochondria, H3 is a maker for nucleus. CD133 is highly enriched in the top fractions.

**Supplementary Figure 5.**
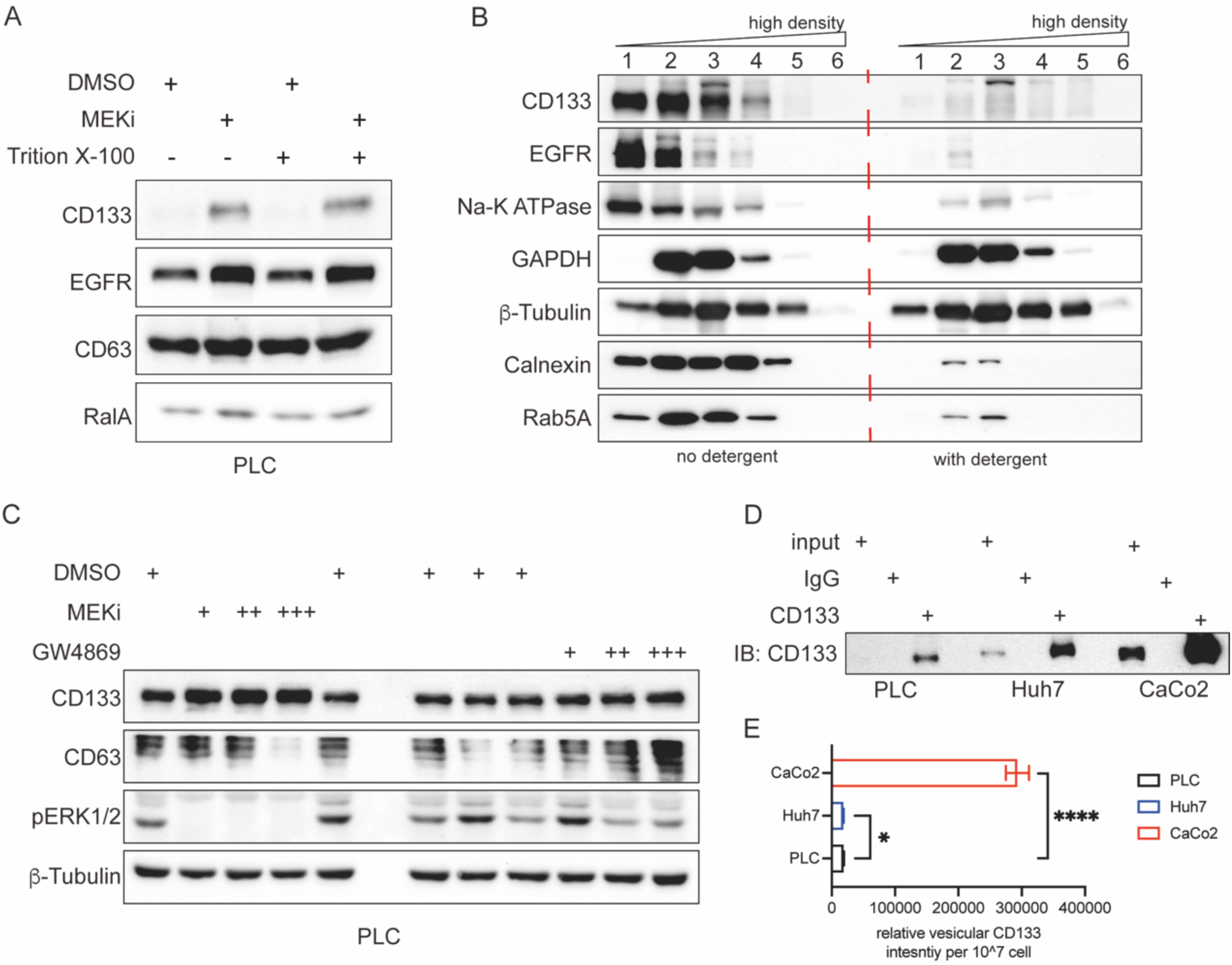
CD133^+^ vesicles are sensitive to detergent and various in different cell lines. (A) Total cell lysates from PLC cells were prepared in the presence or absence from detergent Triton X-100. Then samples were applied to western blot. CD133 protein levels were induced when treated with MEKi no matter of the detergent treatment. (B) Same amount crude vesicles were loaded to high density fractionation after treating with PBS control or 1% Triton X-100. The fractionation results were detected by western blot, the signals of CD133, EGFR, Na^+^- K^+^ ATPase, which were all membrane associated proteins, were significantly decreased in the detergent treated sample. Crude samples were both from MEKi treated PLC cells. (C) Protein level of CD133 was determined by western blot with treatment of MEKi and GW4869 at different concentration in PLC cells. GW4869 is an inhibitor of EV secretion. Blocking EV secretion does not increase intracellular CD133 level, indicating CD133^+^ vesicle is different from EV. (D-E) CD133^+^ vesicles IP from PLC, Huh7 and CaCo2. (D) CD133 protein level detection in CD133^+^ vesicles from different cell lines. (E) Quantification of vesicular CD133 intensity normalized to the cell number. Statistical significance is determined by Student’s t-test. Data were represented as mean ± SEM. *p<0.05, **p<0.01, ***p<0.001.

**Supplementary Figure 6.**
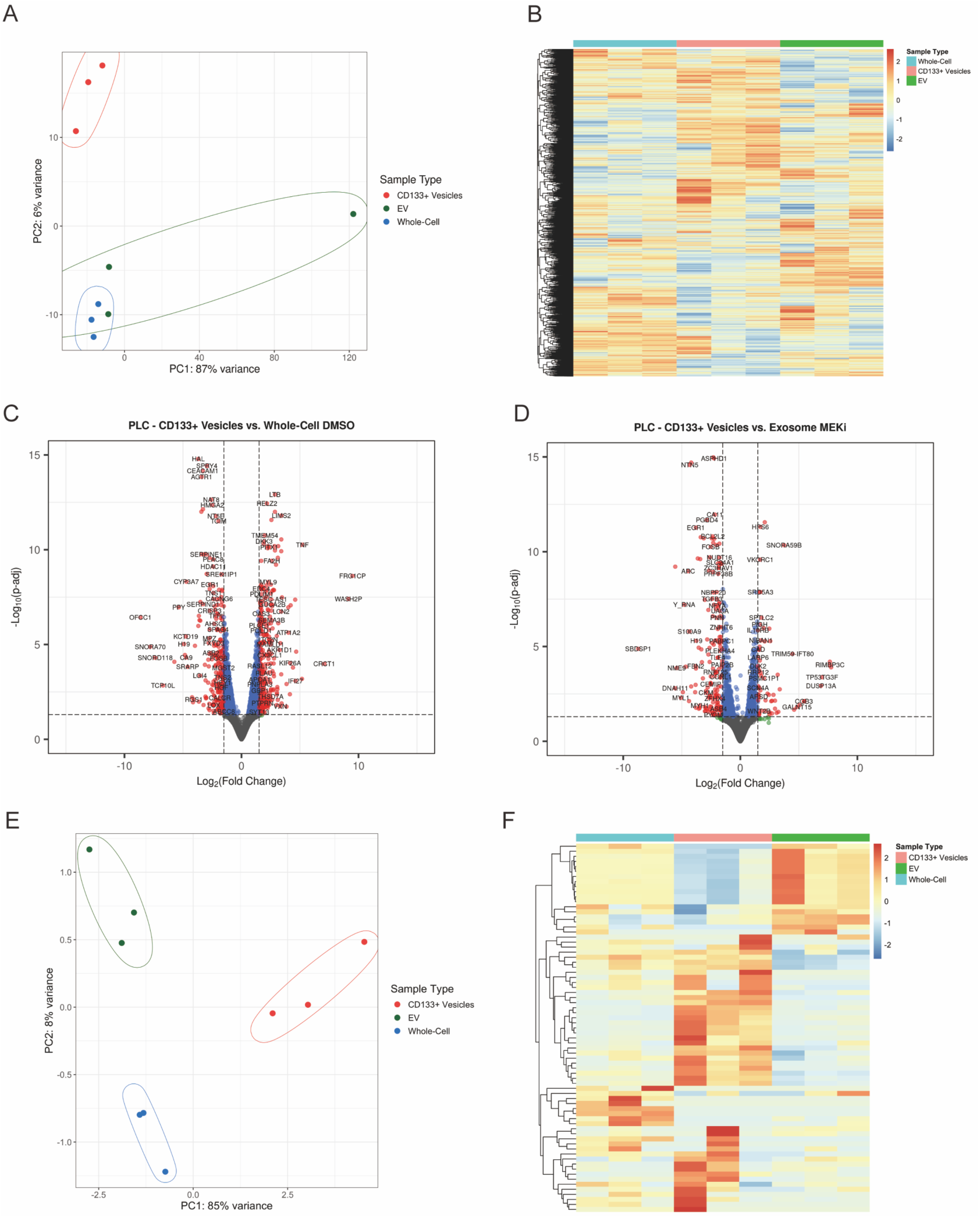
CD133^+^ vesicles have different RNA cargo compared with EVs. (A) Principal Component Analysis (PCA) of gene expression data from whole cell, CD133^+^ vesicles and EVs. Each point represents a biological replicate, with colors denoting the group. Red for CD133^+^ vesicles, Green for EV and Blue for whole cells. (B) A heatmap built with RNA-seq data shows the different gene expression profiles in whole cells, CD133^+^ vesicles and EVs. (C) Volcano plot showing the differential expression of genes between CD133^+^ vesicles and whole cell. The x-axis represents the log_2_ fold change (log_2_FC), with positive values indicating upregulation in the treated group and negative values indicating downregulation. The y-axis shows the −log_10_(p-value), with higher values indicating more statistically significant differences. Points are colored based on p-value significance, with red representing genes that are significantly differentially expressed (p < 0.05) and blue representing non-significant genes. (C) Volcano plot showing the differential expression of genes between CD133^+^ vesicles and EVs. The x-axis represents the log2 fold change (log_2_FC), with positive values indicating upregulation in the treated group and negative values indicating downregulation. The y-axis shows the −log_10_(p-value), with higher values indicating more statistically significant differences. Points are colored based on p-value significance, with red representing genes that are significantly differentially expressed (p < 0.05) and blue representing non- significant genes. (E) Principal Component Analysis (PCA) of small RNA expression data from whole cell, CD133^+^ vesicles and EVs. Each point represents a biological replicate, with colors denoting the group. Red for CD133^+^ vesicles, Green for EV and Blue for whole cells. (F) A heatmap built with small RNA-seq data shows the different small RNA expression profiles in whole cells, CD133^+^ vesicles and EVs.

**Supplementary Figure 7.**
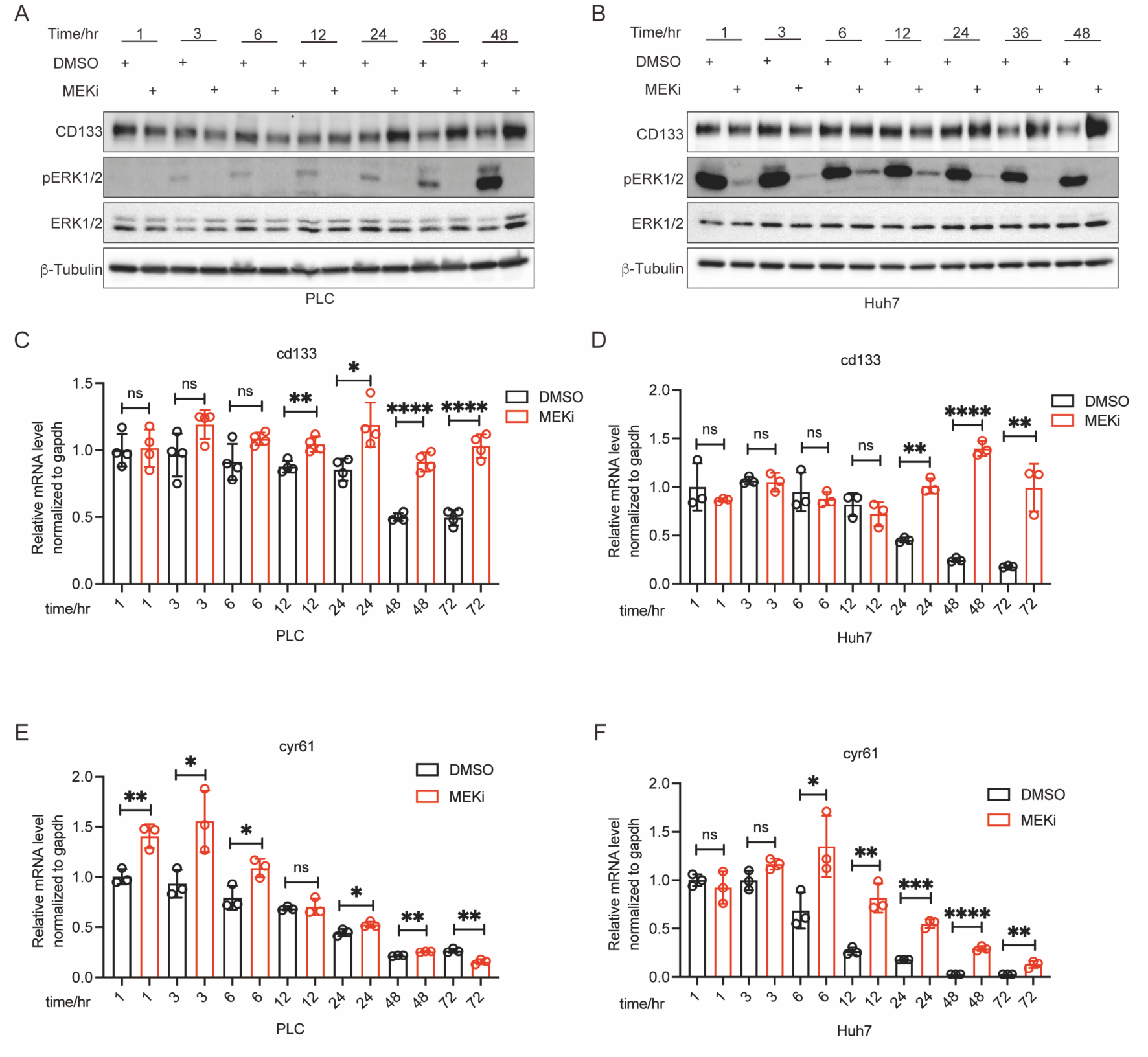
MEK inhibition reveals a coordinated expression pattern between CD133 and Hippo-YAP downstream gene. (A) Protein levels of CD133 and pERK1/2 were analyzed by western blotting at various MEKi treatment time points in PLC cells. (B) Protein levels of CD133 and pERK1/2 were analyzed by western blotting at various MEKi treatment time points in Huh7 cells. (C) qRT-PCR analysis of CD133 mRNA levels in PLC cells treated with MEKi over time. PLC cells were treated with MEKi for varying durations (e.g., 1, 3, 6, 12, 24, 48 and 72 hours). (D) qRT-PCR analysis of CYR61 mRNA levels in PLC cells treated with MEKi over time. PLC cells were treated with MEKi for varying durations (e.g., 1, 3, 6, 12, 24, 48 and 72 hours). (E) qRT-PCR analysis of CD133 mRNA levels in Huh7 cells treated with MEKi over time. Huh7 cells were treated with MEKi for varying durations (e.g., 1, 3, 6, 12, 24, 48 and 72 hours). (F) qRT-PCR analysis of CYR61 mRNA levels in Huh7 cells treated with MEKi over time. Huh7 cells were treated with MEKi for varying durations (e.g., 1, 3, 6, 12, 24, 48 and 72 hours). Statistical significance is determined by Student’s t-test. Data were represented as mean ± SEM. *p<0.05, **p<0.01, ***p<0.001, ns: no significance.

**Supplementary Figure 8.**
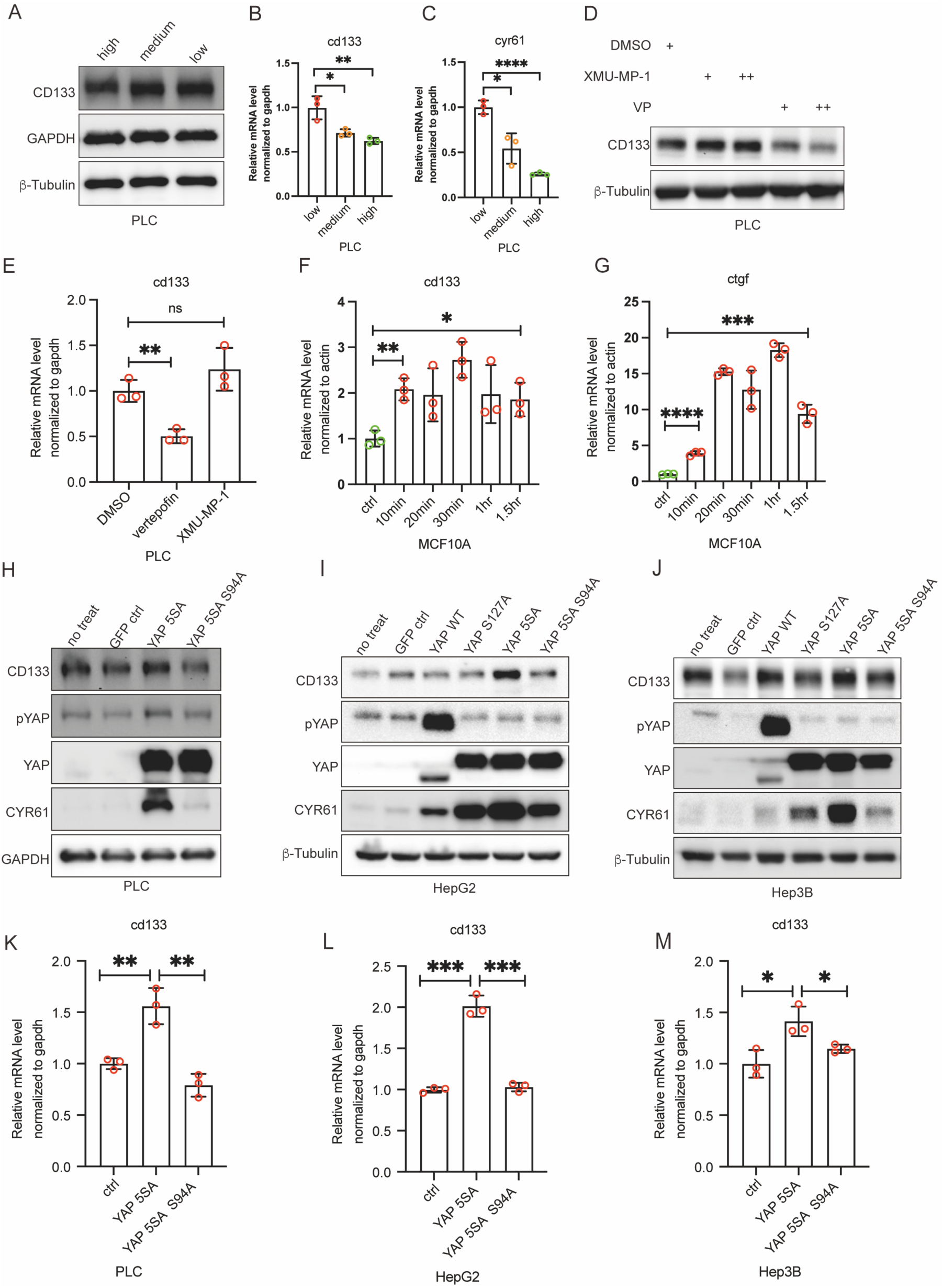
CD133 is regulated by Hippo-YAP signal pathway. (A) Protein level of CD133 was analyzed by western blotting at different cell density in PLC cells. (B-C) mRNA levels of CD133 (B) and cyr61 (C) were analyzed by qRT-PCR at different cell density in PLC cells. (D) Protein level of CD133 was analyzed by western blotting when treating PLC cells with XMU-MP-1 and VP. (E) mRNA level of CD133 was analyzed by qRT-PCR when treating PLC cells with XMU-MP-1 and VP. (F-G) mRNA level of CD133 and ctgf were analyzed by qRT-PCR in the serum stimulation experiments. CD133 and ctgf were upregulated in response to serum recovery in MCF10A cells. MCF10A cells were starved in SFM for 12 hours and then cells were treated with 10% FBS for the indicated time. (F) CD133 mRNA level, (G) ctgf mRNA level. (H) Protein level of CD133 was analyzed by western blotting in YAP-5SA and YAP-5SA/S94A overexpression PLC cells. (I) Protein level of CD133 was analyzed by western blotting in YAP-WT, YAP-S127A, YAP-5SA and YAP- 5SA/S94A overexpression HepG2 cells. (J) Protein level of CD133 was analyzed by western blotting in YAP-WT, YAP-S127A, YAP-5SA and YAP- 5SA/S94A overexpression Hep3B cells. (K) mRNA level of CD133 was analyzed in YAP-5SA overexpression and YAP-5SA/S94A overexpression PLC cells. (L) mRNA level of CD133 was analyzed in YAP-5SA overexpression and YAP-5SA/S94A overexpression HepG2 cells. (M) mRNA level of CD133 was analyzed in YAP-5SA overexpression and YAP-5SA/S94A overexpression Hep3B cells. Statistical significance is determined by Student’s t-test. Data were represented as mean ± SEM. *p<0.05, **p<0.01, ***p<0.001, ns: no significance.

## References

1. Hanahan D. Hallmarks of Cancer: New Dimensions. Cancer Discov 2022;12:31–46.

2. Holohan C, Van Schaeybroeck S, Longley DB, Johnston PG. Cancer drug resistance: an evolving paradigm. Nat Rev Cancer 2013;13:714–726.

3. Vasan N, Baselga J, Hyman DM. A view on drug resistance in cancer. Nature 2019;575:299–309.

4. Lavoie H, Gagnon J, Therrien M. ERK signalling: a master regulator of cell behaviour, life and fate. Nat Rev Mol Cell Biol 2020;21:607–632.

5. Scheiter A, Lu LC, Gao LH, Feng GS. Complex Roles of PTPN11/SHP2 in Carcinogenesis and Prospect of Targeting SHP2 in Cancer Therapy. Annu Rev Cancer Biol 2024;8:15–33.

6. Sodir NM, Pathria G, Adamkewicz JI, Kelley EH, Sudhamsu J, Merchant M, Chiarle R, et al. SHP2: A Pleiotropic Target at the Interface of Cancer and Its Microenvironment. Cancer Discov 2023;13:2339–2355.

7. Kaneko K, Liang Y, Liu Q, Zhang S, Scheiter A, Song D, Feng GS. Identification of CD133(+) intercellsomes in intercellular communication to offset intracellular signal deficit. Elife 2023;12.

8. Bard-Chapeau EA, Li S, Ding J, Zhang SS, Zhu HH, Princen F, Fang DD, et al. Ptpn11/Shp2 acts as a tumor suppressor in hepatocellular carcinogenesis. Cancer Cell 2011;19:629–639.

9. Marzesco AM. Prominin-1-containing membrane vesicles: origins, formation, and utility. Adv Exp Med Biol 2013;777:41–54.

10. Izumi H, Li Y, Shibaki M, Mori D, Yasunami M, Sato S, Matsunaga H, et al. Recycling endosomal CD133 functions as an inhibitor of autophagy at the pericentrosomal region. Sci Rep 2019;9:2236.

11. Oh HT, Heo W, Yoo GD, Kim KM, Hwang JH, Hwang ES, Ko J, et al. CD133-Src-TAZ signaling stimulates ductal fibrosis following DDC diet-induced liver injury. J Cell Physiol 2022;237:4504–4516.

12. Lee H, Yu DM, Bahn MS, Kwon YJ, Um MJ, Yoon SY, Kim KT, et al. Hepatocyte-specific Prominin-1 protects against liver injury-induced fibrosis by stabilizing SMAD7. Exp Mol Med 2022;54:1277–1289.

13. You H, Ding W, Rountree CB. Epigenetic regulation of cancer stem cell marker CD133 by transforming growth factor-beta. Hepatology 2010;51:1635–1644.

14. Wei Y, Jiang Y, Zou F, Liu Y, Wang S, Xu N, Xu W, et al. Activation of PI3K/Akt pathway by CD133-p85 interaction promotes tumorigenic capacity of glioma stem cells. Proc Natl Acad Sci U S A 2013;110:6829–6834.

15. Bahn MS, Yu DM, Lee M, Jo SJ, Lee JW, Kim HC, Lee H, et al. Central role of Prominin- 1 in lipid rafts during liver regeneration. Nat Commun 2022;13:6219.

16. Moreno-Londono AP, Robles-Flores M. Functional Roles of CD133: More than Stemness Associated Factor Regulated by the Microenvironment. Stem Cell Rev Rep 2024;20:25–51.

17. Behrooz AB, Syahir A, Ahmad S. CD133: beyond a cancer stem cell biomarker. Journal of Drug Targeting 2019;27:257–269.

18. Glumac PM, LeBeau AM. The role of CD133 in cancer: a concise review. Clinical and Translational Medicine 2018;7.

19. Soleimani A, Dadjoo P, Avan A, Soleimanpour S, Rajabian M, Ferns G, Ryzhikov M, et al. Emerging roles of CD133 in the treatment of gastric cancer, a novel stem cell biomarker and beyond. Life Sciences 2022;293.

20. Yin S, Li J, Hu C, Chen X, Yao M, Yan M, Jiang G, et al. CD133 positive hepatocellular carcinoma cells possess high capacity for tumorigenicity. Int J Cancer 2007;120:1444–1450.

21. Yang K, Zhao Y, Du Y, Tang R. Evaluation of Hippo Pathway and CD133 in Radiation Resistance in Small-Cell Lung Cancer. J Oncol 2021;2021:8842554.

22. Lee TK, Guan XY, Ma S. Cancer stem cells in hepatocellular carcinoma - from origin to clinical implications. Nat Rev Gastroenterol Hepatol 2022;19:26–44.

23. Loh JJ, Ma S. Hallmarks of cancer stemness. Cell Stem Cell 2024;31:617–639.

24. Franklin JM, Wu Z, Guan KL. Insights into recent findings and clinical application of YAP and TAZ in cancer. Nat Rev Cancer 2023;23:512–525.

25. Feng X, Lu T, Li J, Yang R, Hu L, Ye Y, Mao F, et al. The Tumor Suppressor Interferon Regulatory Factor 2 Binding Protein 2 Regulates Hippo Pathway in Liver Cancer by a Feedback Loop in Mice. Hepatology 2020;71:1988–2004.

26. Johnson R, Halder G. The two faces of Hippo: targeting the Hippo pathway for regenerative medicine and cancer treatment. Nat Rev Drug Discov 2014;13:63–79.

27. Moya IM, Halder G. Hippo-YAP/TAZ signalling in organ regeneration and regenerative medicine. Nat Rev Mol Cell Biol 2019;20:211–226.

28. Russell JO, Camargo FD. Hippo signalling in the liver: role in development, regeneration and disease. Nat Rev Gastroenterol Hepatol 2022;19:297–312.

29. Zanconato F, Cordenonsi M, Piccolo S. YAP/TAZ at the Roots of Cancer. Cancer Cell 2016;29:783–803.

30. Zhu N, Yang R, Wang X, Yuan L, Li X, Wei F, Zhang L. The Hippo signaling pathway: from multiple signals to the hallmarks of cancers. Acta Biochim Biophys Sin (Shanghai) 2023;55:904–913.

31. Feng X, Wang Z, Wang F, Lu T, Xu J, Ma X, Li J, et al. Dual function of VGLL4 in muscle regeneration. EMBO J 2019;38:e101051.

32. Jeppesen DK, Fenix AM, Franklin JL, Higginbotham JN, Zhang Q, Zimmerman LJ, Liebler DC, et al. Reassessment of Exosome Composition. Cell 2019;177:428–445 e418.

33. Cao M, Isaac R, Yan W, Ruan X, Jiang L, Wan Y, Wang J, et al. Cancer-cell-secreted extracellular vesicles suppress insulin secretion through miR-122 to impair systemic glucose homeostasis and contribute to tumour growth. Nat Cell Biol 2022;24:954–967.

34. Kalluri R, LeBleu VS. The biology, function, and biomedical applications of exosomes. Science 2020;367.

35. Qi S, Zhong Z, Zhu Y, Wang Y, Ma M, Wang Y, Liu X, et al. Two Hippo signaling modules orchestrate liver size and tumorigenesis. EMBO J 2023;42:e115749.

36. Guo X, Zhao Y, Yan H, Yang Y, Shen S, Dai X, Ji X, et al. Single tumor-initiating cells evade immune clearance by recruiting type II macrophages. Genes Dev 2017;31:247–259.

37. Zhang S, Pan C, Lv X, Wu W, Chen H, Wu W, Wu H, et al. Repression of Abd-B by Polycomb is critical for cell identity maintenance in adult Drosophila testis. Sci Rep 2017;7:5101.

38. Zhang S, Zhao J, Lv X, Fan J, Lu Y, Zeng T, Wu H, et al. Analysis on gene modular network reveals morphogen-directed development robustness in Drosophila. Cell Discov 2020;6:43.

